# Mercury methylation by metabolically versatile and cosmopolitan marine bacteria

**DOI:** 10.1101/2020.06.03.132969

**Authors:** Heyu Lin, David B. Ascher, Yoochan Myung, Carl H. Lamborg, Steven J. Hallam, Caitlin M. Gionfriddo, Kathryn E. Holt, John W. Moreau

## Abstract

Microbes transform aqueous mercury (Hg) into methylmercury (MeHg), a potent neurotoxin in terrestrial and marine food webs. This process requires the gene pair *hgcAB*, which encodes for proteins that actuate Hg methylation, and has been well described for anoxic environments. However, recent studies report potential MeHg formation in suboxic seawater, although the microorganisms involved remain poorly understood. In this study, we conducted large-scale multi-omic analyses to search for putative microbial Hg methylators along defined redox gradients in Saanich Inlet (SI), British Columbia, a model natural ecosystem with previously measured Hg and MeHg concentration profiles. Analysis of gene expression profiles along the redoxcline identified several putative Hg methylating microbial groups, including *Calditrichaeota*, SAR324 and *Marinimicrobia*, with the last by far the most active based on *hgc* transcription levels. *Marinimicrobia hgc* genes were identified from multiple publicly available marine metagenomes, consistent with a potential key role in marine Hg methylation. Computational homology modelling predicted that *Marinimicrobia* HgcAB proteins contain the highly conserved structures required for functional Hg methylation. Furthermore, a number of terminal oxidases from aerobic respiratory chains were associated with several SI putative novel Hg methylators. Our findings thus reveal potential novel marine Hg-methylating microorganisms with a greater oxygen tolerance and broader habitat range than previously recognised.

## Introduction

Mercury (Hg), a highly toxic metal, is widespread in the environment from primarily anthropogenic sources, leading to increased public concern over the past few decades. (e.g. Fitzgerald and Clarkson 1991, Hsu-Kim et al 2018, Selin 2009). Methylmercury (MeHg) is recognized as a potent neurotoxin that bioaccumulates through both marine and terrestrial food webs (Lee and Fisher 2017, Selin 2009). With implementation of the Minamata Convention on Mercury (Hsu-Kim et al 2018), better understanding is expected of potential MeHg sources, in the context of global biogeochemical cycles and factors influencing Hg speciation. For example, oxygen gradients in seawater have been observed to expand in response to climate change (Stramma et al 2008), and the resulting impacts on biogeochemical cycles, e.g., carbon, nitrogen, and sulfur, have been studied (e.g., Wright et al 2012). Effects on Hg cycling, however, are rarely considered.

The environmental transformation of Hg(II) to MeHg is a microbially mediated process, for which the proteins are encoded by the two-gene cluster *hgcA* and *hgcB* (Parks et al 2013). Possession of the *hgcAB* gene pair is a predictor for Hg methylation capability (Gilmour et al 2013), and the discovery of *hgc* has stimulated a search for potential Hg methylating microbes in diverse environments (Podar et al 2015). To date, all experimentally confirmed Hg methylators are anaerobes (Grégoire et al 2018, Podar et al 2015) from: three *Deltaproteobacteria* clades (sulfate-reducing bacteria, SRB; Fe-reducing bacteria, FeRB); and syntrophic bacteria *Syntrophobacterales*), one clade belonging to fermentative *Firmicutes*, and an archaeal clade, *Methanomicrobia*. These microorganisms are ubiquitous in soils, sediments, seawater, freshwater, and extreme environments, as well as the digestive tracts of some animals (Podar et al 2015). Recently, other *hgc* carriers, not only from anaerobic but also microaerobic habitats, were discovered using culture-independent approaches, including *Chloroflexi, Chrysiogenetes, Spirochaetes* (Podar et al 2015), *Nitrospina* (Gionfriddo et al 2016), and *Verrucomicrobia* (Jones et al 2019).

A global Hg survey (Lamborg et al 2014) found a prevalence for MeHg in suboxic waters, especially in regionally widespread oxygen gradients at upper and intermediate ocean depths. Oxygen concentrations are lower there due to a combination of physical and biological forcing effects (Paulmier and Ruiz-Pino 2009, Wright et al 2012). These gradients can exist as permanent features in the water column, impinge on coastal margins, or manifest more transiently (e.g., induced by phytoplankton blooms). As oxygen levels decrease, metabolic energy gets increasingly diverted to alternative electron acceptors, resulting in coupling of other biogeochemical cycles, e.g., C, N, S, Fe, and Mn (Bertagnolli and Stewart 2018, Moore and Doney 2007, Wright et al 2012). Recent findings of microaerophilic microbial Hg methylation potential in sea ice and seawater (Gionfriddo et al 2016, Villar et al 2020) raise the possibility that this process contributes significantly to ocean MeHg biomagnification.

In this study, we used existing Hg (total) and MeHg concentration data from Saanich Inlet (SI), a seasonally anoxic fjord on the coast of Vancouver Island (British Columbia, Canada) to guide targeted metagenomic and metatranscriptomic analyses of SI seawater from varying depths. Saanich Inlet, as a model natural ecosystem for studying microbial activity along defined oxygen gradients (Hawley et al 2017b), provides an ideal site to search for novel putative Hg methylators in low-oxygen environments. Computational homology modelling was performed to predict the functionality of HgcAB proteins encoded for by putative novel Hg-methylators. Finally, we scanned global metagenomic datasets for recognizable *hgcA* genes, to reassess the environmental distribution of microbial mercury methylation potential.

## Results and Discussion

### Hg and MeHg concentrations along redox gradients in SI

Concentrations of total dissolved Hg (Hg_T_) and monomethylmercury (MeHg) from filtered SI station “S3” seawater samples were vertical profiled from eight different depths (10 m to 200 m) below sea surface. The concentration of Hg_T_ at sea surface was ∼0.70 pM and remained nearly constant in seawater above 120 m depth, increasing to 1.35 pM and ∼10.56 pM at 135 m and 200 m depths, respectively (Figure 1A). MeHg was below detection limit (<0.1 pM) for seawater above 100 m depth, but increased to 0.50 pM (17.2% of total Hg) at 150 m depth. However, MeHg then decreased to 0.1 pM at 165 m depth, becoming undetectable at 200 m (Figure 1B and C).

**Figure 1.**
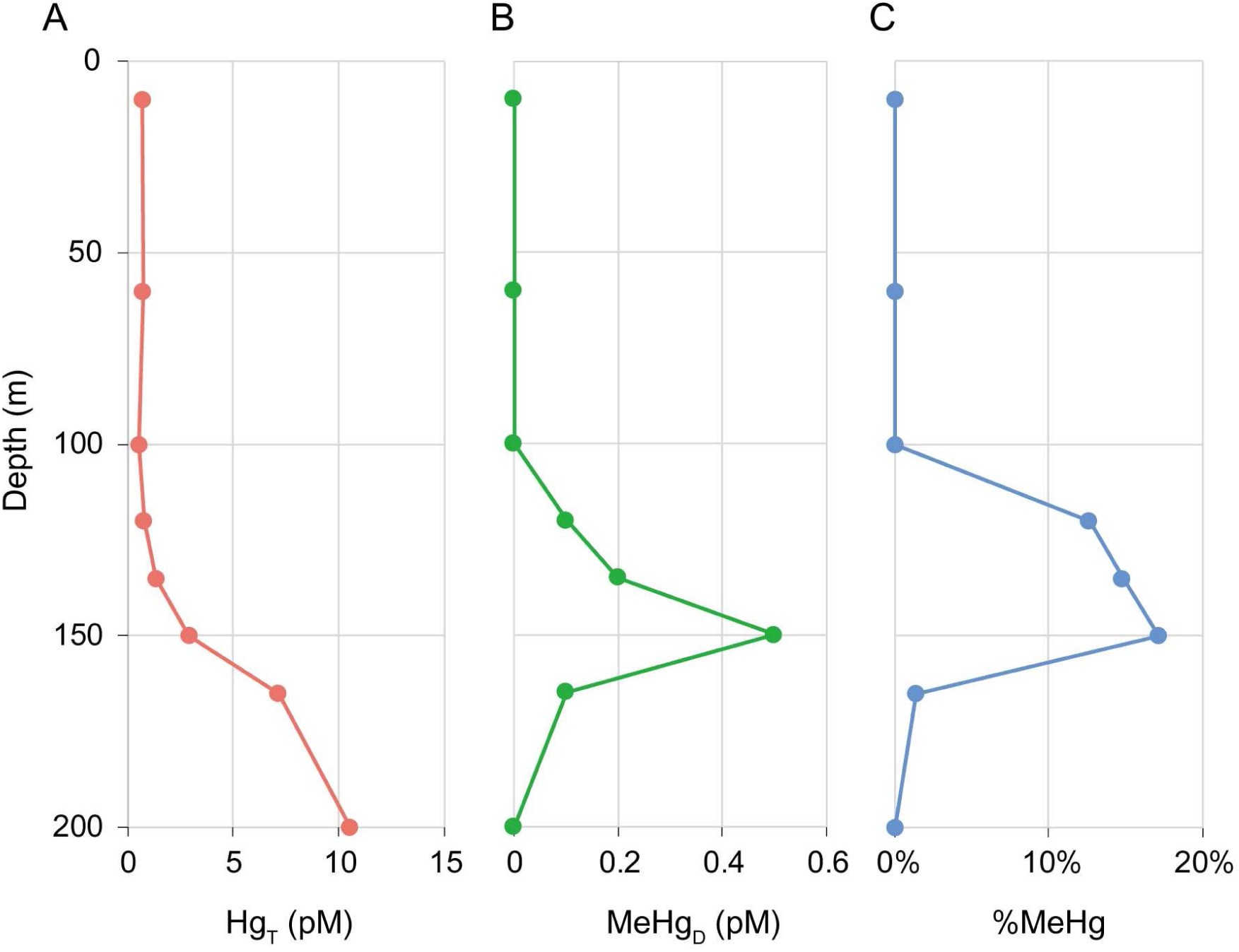
Hg and MeHg profiles of Saanich Inlet station “S3”. (A) Concentration of total dissolved Hg. (B) Concentration of dissolved MeHg. (C) MeHg as a percentage of total dissolved Hg.

### MAG (metagenome-assembled genome) reconstruction and putative Hg methylator identification

A total of 2088 MAGs with completeness >70% and contamination <5% were recovered from all SI metagenomic datasets (generated across all sampling depths, 96% being from station S3). From these, 56 MAGs, belonging to seven phyla: *Proteobacteria, Marinimicrobia, Verrucomicrobia, Firmicutes, Calditrichaeota, Spirochaetes*, and *Nitrospinae* (Table S1), were identified as having genes homologous with *hgcA* (Figure S1). Among these, *Marinimicrobia* and *Calditrichaeota* have not previously been implicated in Hg methylation. No *hgcAB* genes were recovered from archaeal MAGs, suggesting that archaeal Hg methylators were rare or absent.

### Novel potential Hg methylators

Fifteen *hgcA*-carrying MAGs represented members of phylum *Marinimicrobia*, a widespread but uncultured marine phylum that couples C, N, and S cycling (Hawley et al 2017a). These *Marinimicrobia* contained the same *hgcA* gene sequence (100% nucleotide identity), and uniformly possessed *hgcB* genes downstream of *hgcA. Marinimicrobia* genome association with *hgcAB* was supported strongly by emergent self-organizing maps (ESOMs; Figure S2A). One *Marinimicrobia-*associated MAG, SI037_bin139, exhibited the highest binning quality score (Table S1), with a completeness of 97.8% and contamination ∼0%, having 75 contigs in total. Many *Marinimicrobia* MAGs also carried 16S rRNA genes, further supporting taxonomic classification (Figure S3). Notably, *Marinimicrobia*-HgcA sequences fell within the *Euryarchaeota* in the HgcA phylogenetic tree (Figure 2), suggesting a different evolutionary pathway to other bacterial *hgc* sequences.

**Figure 2.**
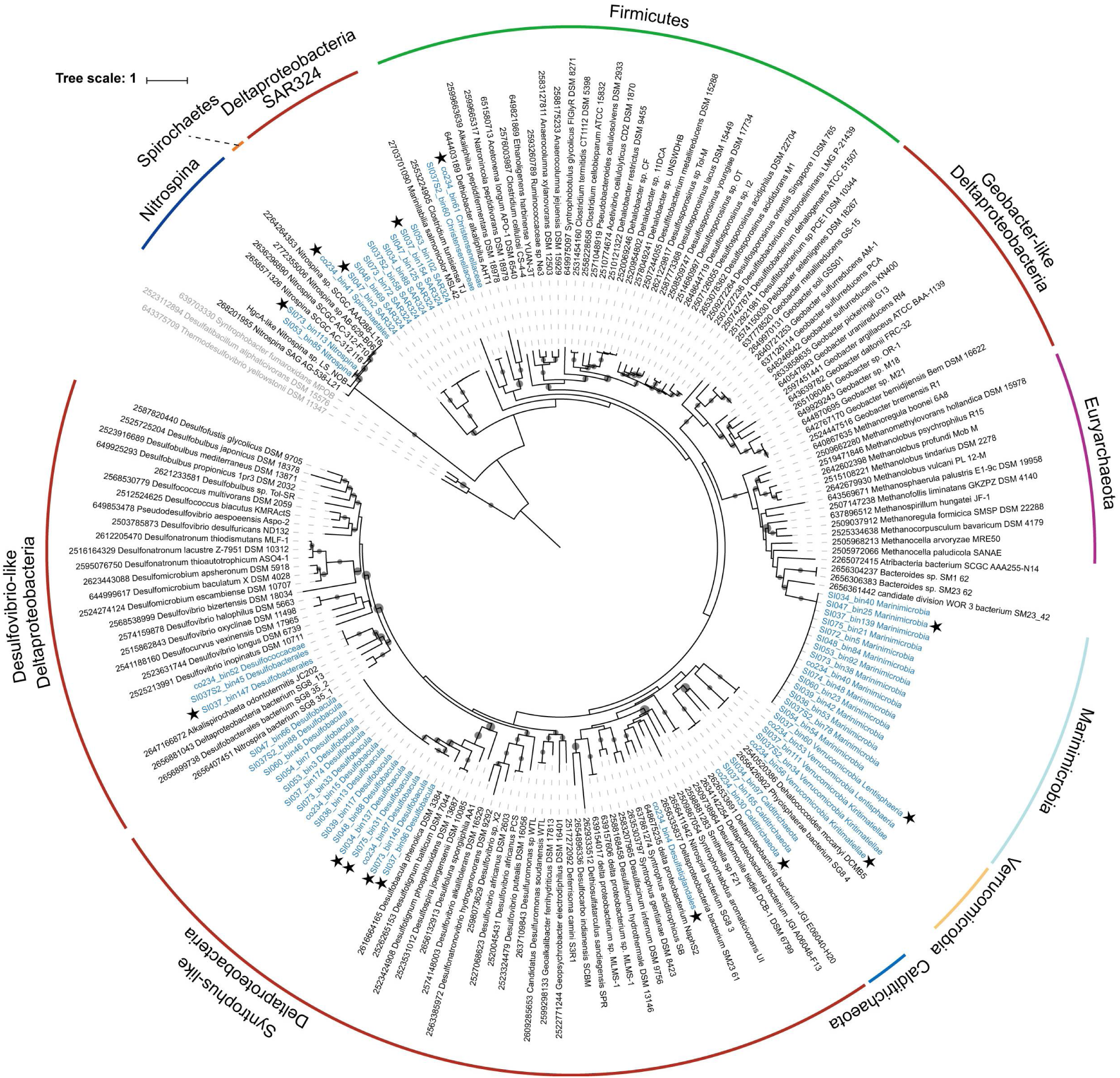
Maximum-likelihood phylogenetic tree of HgcA amino acid sequences. (1,000 ultrafast bootstrap replicates; values >90% are shown by black dots at the nodes). HgcA sequences recovered in this study are highlighted in blue. HgcA sequences retrieved from public databases with corresponding IMG gene IDs are shown in black. HgcA paralogues from non-methylating bacteria were used as outgroups and are shown in grey. The 15 representative HgcA sequences used in this study are indicated by stars. Taxonomic classifications of the *hgcA*-carrying genomes are labeled in the outer circle by different colors.

Three MAGs containing *hgcAB* genes were associated with *Calditrichaeota*, another cosmopolitan marine phylum (Marshall et al 2017). HgcA sequences from *Calditrichaeota* formed a cluster with the *Planctomycetes, Verrucomicrobia* and *Chlamydiae* (PVC) group and *Chloroflexi*, which also have close phylogenetic relationships to *Deltaproteobacteria*, as shown in Figure 2. ESOMs (Figure S2B) supported the presence of *hgcAB* genes in *Calditrichaeota* MAGs.

HgcA sequences were found in eight *Proteobacteria* MAGs, all belonging to the *Deltaproteobacteria* group SAR324. *Deltaproteobacteria* are the most diverse Hg-methylating clade currently known (Podar et al 2015). However, the SAR324 *hgcA* sequences clustered separately from those of three previously described *Deltaproteobacteria* methylating clades (*Desulfovibrio*-like, *Geobacter*-like, and *Syntrophus*-like; Gilmour et al, 2013; Podar et al, 2015), forming a potentially novel clade of Hg-methylators (Figure 2). SAR324 is affiliated with a group of *Deltaproteobacteria* abundant in the deep sea and in low-oxygen settings, with the ability to metabolize sulfur, organic carbon, and C_1_ compounds (Sheik et al 2014). An adjacent *hgcB* gene was also recognised downstream of *hgcA* in SAR324 MAGs SI047_bin2 and SI048_bin69. Six other SAR324 MAGs did not contain recognizable *hgcAB* genes, but several HgcB-like candidates with tandem [CX2CX2CX3C] motifs were recognised. The presence of *hgc* genes in SAR324 MAGs was well supported by ESOM (Figure S2A).

### Relative abundance of hgcA DNA and RNA in SI

The 56 *hgcA* sequences were clustered into 15 groups using a 99% sequence identity threshold. One *hgcA* sequence was selected arbitrarily from each group, with corresponding MAG (Table 1; also indicated by stars in Figure 2), for further analysis. These sequences were used to recruit reads from SI metagenomic and metatranscriptomic datasets, in order to calculate relative abundance and expression. Results showed that *hgcA* genes were widely distributed in SI water samples, and the abundance of *hgcA* gene copies increased with depth, peaking at ∼200 m depth (Figure 3A). Since *hgcA* is usually present as a single copy per genome, this finding suggests that Hg methylators were most abundant at this depth. Notably, *Marinimicrobia* was the most abundant putative Hg-methylator detected throughout the water column, accounting for >2% of the microbial community for many samples from 200 m depth (Figure 3B). *Deltaproteobacteria* with *hgcA* also showed higher abundance in 200 m depth samples, suggesting an overlapping habitat range with *Marinimicrobia*. Although *hgcA*-carrying *Nitrospina* was not a dominant phylum in SI datasets, the relative abundance of *Nitrospina-*hosted *hgcA* remained nearly constant from sea surface to bottom water, implying the adaptability this phylum across the redoxcline.

**Table 1.** Summary of 15 representative *hgcA*-carrying MAGs ^a^.

| MAGs | %Compl.<sup>b</sup> | %Conta.<sup>b</sup> | Size (Mbp) | # Contigs | %GC | 16S | Taxonomy |
| --- | --- | --- | --- | --- | --- | --- | --- |
| SI034_bin137 | **97.58** | **2.992** | 4.94 | 304 | 40.0 |  | *Desulfobacula sp.* |
| SI037_bin96 | 71.02 | **3.87** | 2.76 | 633 | 39.5 |  | *Desulfobacula sp.* |
| SI073_bin145 | 87.48 | 5.322 | 4.26 | 657 | 39.2 |  | *Desulfobacula sp.* |
| SI075_bin31 | 88.61 | **2.293** | 4.89 | 394 | 40.1 |  | *Desulfobacula sp.* |
| co234_bin4 | 81.15 | 5.545 | 5.91 | 1000 | 50.1 |  | Order *Desulfatiglandales* |
| SI037_bin147 | 89.67 | **1.935** | 3.28 | 289 | 38.6 |  | Order *Desulfobacterales* |
| SI037_bin154 | 89.02 | **3.361** | 7.85 | 512 | 45.5 |  | SAR324 group |
| SI047_bin2 | 74.15 | **2.941** | 3.63 | 142 | 36.2 |  | SAR324 group |
| SI047_bin25 | **96.7** | **0** | 3.17 | 92 | 37.4 | + | Phylum *Marinimicrobia* |
| co234_bin30 | **96.08** | **1.648** | 4.54 | 293 | 42.9 |  | Phylum *Calditrichaeota* |
| SI037_bin60 | 83.67 | **3.914** | 8.81 | 2073 | 63.9 |  | Class *Lentisphaeria* |
| co234_bin56 | **95.6** | **2.027** | 4.85 | 162 | 60.6 | + | Class *Kiritimatiellae* |
| co234_bin61 | **98.25** | **0** | 2.67 | 97 | 47.7 |  | Order *Christensenellales* |
| co234_bin41 | 72.27 | **1.253** | 4.68 | 1025 | 48.9 |  | Order *Spirochaetales* |
| SI073_bin113 | 74.89 | **1.709** | 2.92 | 321 | 43.1 |  | *Nitrospina sp.* |

<sup>a</sup> Representative MAGs selected by 99% *hgcA* identity threshold. See Table S1 for all 56 *hgcA*-carrying MAGs.
<sup>b</sup> Completeness above 95% and contamination below 5% are shown in bold; quality of MAGs was determined by CheckM with the “lineage\_wf” pipeline.

**Figure 3.**
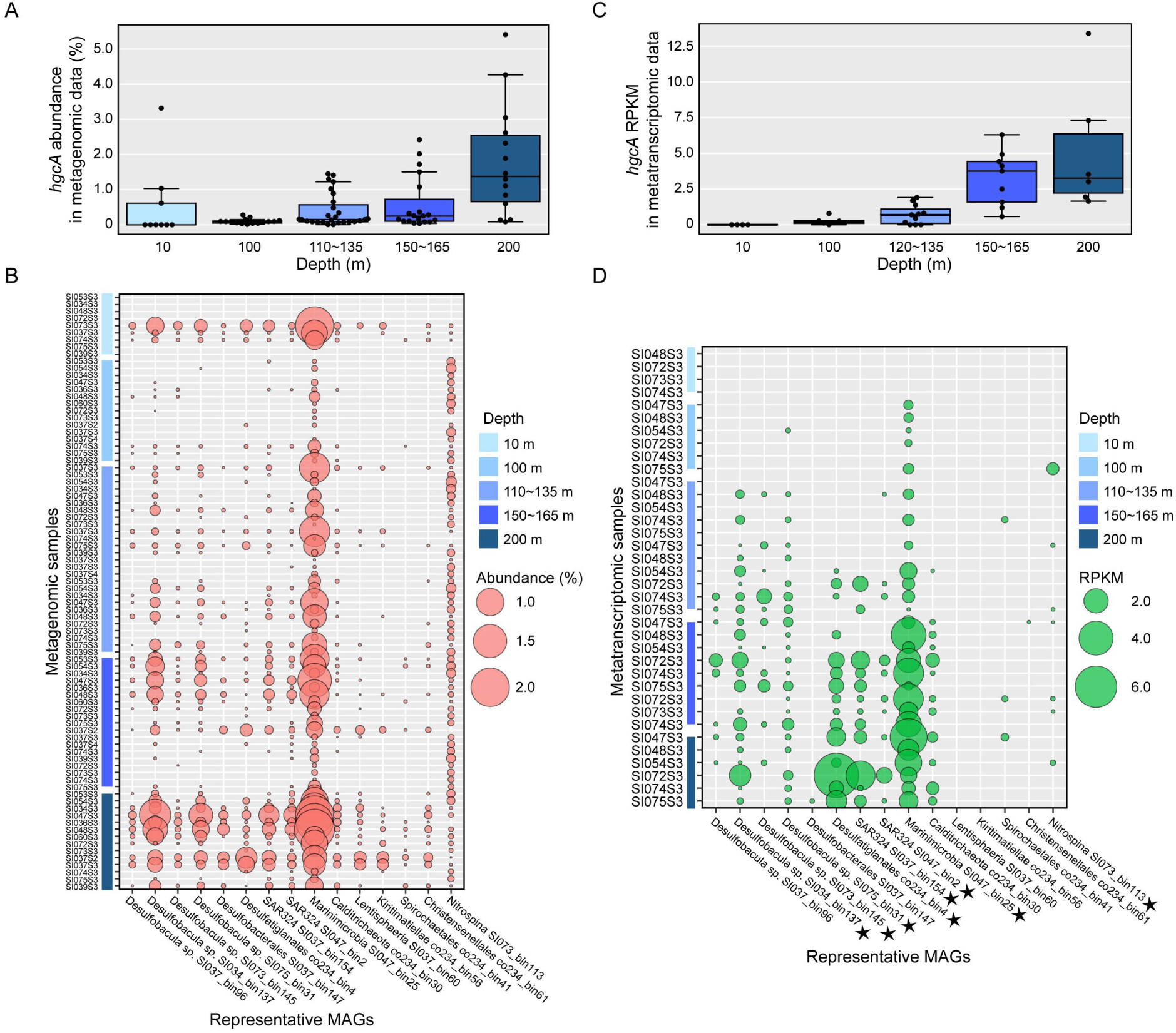
Relative abundance of representative *hgcA* genes and transcripts in different samples from Saanich Inlet. Gene abundance is normalized by gene length and genome equivalent for each sample, and transcript abundances are represented as RPKM values. (A) Relative abundance of the sum of representative *hgcA* genes for different depths. (B) Relative abundance of different representative *hgcA* genes from different metagenomic datasets. Larger circles indicate a higher percentage of the whole microbial community. Different shades of blue indicate different depths from which the samples were taken. (C) RPKM values of the sum of representative *hgcA* transcripts in different depths. (D) RPKM values of different representative *hgcA* transcripts in different metatranscriptomic samples. HgcA sequences from metaproteomic samples are indicated with stars.

Reads per kilobase per million mapped (RPKM) values were calculated for *hgcA* transcripts in each sample to assess *hgc* gene expression in a subset of 36 samples from site S3 (Figure 3C). Consistent with relative DNA abundance, *hgcA* transcripts also increased with depth to a maximum at 200 m (mean RPKM = 5.1); the most abundant *hgcA* transcripts were associated with *Marinimicrobia* (Figure 3D). Intriguingly, no *hgcA* transcripts were recovered from 10 m depth samples, although a number of *hgc* DNA sequences were recovered. Additionally, no *hgcA* transcript associated with *Verrucomicrobia* (*Lentisphaeria* and *Kiritimatiellae*) was detected for any dataset, which may reflect the relatively lower abundance of *Verrucomicrobia-hgcA* genes, as well as a low transcription level. Predicted amino acid sequences from SI metaproteomic datasets were also scanned for expression of putative *hgcA* genes (Figure S4; also indicated by stars in Figure 3D). Excepting from phyla *Calditrichaeota, Verrucomicrobia, Spirochaetes*, and *Firmicutes*, representative HgcA sequences could be detected in matched metaproteomic datasets.

### Abundance of merB genes

As MeHg concentrations in the SI water column may reflect net Hg methylation and demethylation, the distribution with depth of *merB*, a gene encoding for an organomercury lyase, was also assessed (Figure 1B). Results showed the relative abundance of *merB* genes peaked at 200 m depth, as ∼0.6% of total microbial community genes (Figure S5A). RPKM values of *merB* transcripts in each metatranscriptomic sample were calculated and compared with *hgcA* transcripts. Consistent with metagenomic results, *merB* transcripts peaked at 200 m depth (mean RPKM = 5.2), whereas no *merB* transcripts were detected at 10 m depth (Figure S5B). Average RPKM values of *merB* were higher than those of *hgcA* at depths < 120 m, but were exceeded by *hgcA* RPKM values at 120-200 m depth. Interestingly, the average RPKM of *hgcA* was again exceeded by that of *merB* at 200 m depth, although transcription of both genes increased to maxima at that same depth (Figure S5C).

### Phylogenetic analysis of hgcA-carrying Marinimicrobia

All 15 *hgcA*-carrying *Marinimicrobia* and 409 non-*hgcA*-carrying *Marinimicrobia* spp., derived from this study and public databases (NCBI and IMG), were used to build a phylogenetic tree based on concatenated alignment of housekeeping genes (Figure 4A, Figure S6). All 15 *hgcA*-carrying *Marinimicrobia* clustered into a monophyletic clade (Figure 4A) and carried identical *hgcA* gene sequences (Figure 2), consistent with a single horizontal transfer of *hgcA* into a sublineage of *Marinimicrobia*. Furthermore, while all *Marinimicrobia*-associated *hgcA* sequences were identical, four different 16S rRNA genes were represented by these MAGs, with minimum identity of 96.0% (Figure S3). This finding suggests that *hgcA*-carrying *Marinimicrobia* represent a novel candidate species or genus, according to Yarza et al (2014).

**Figure 4.**
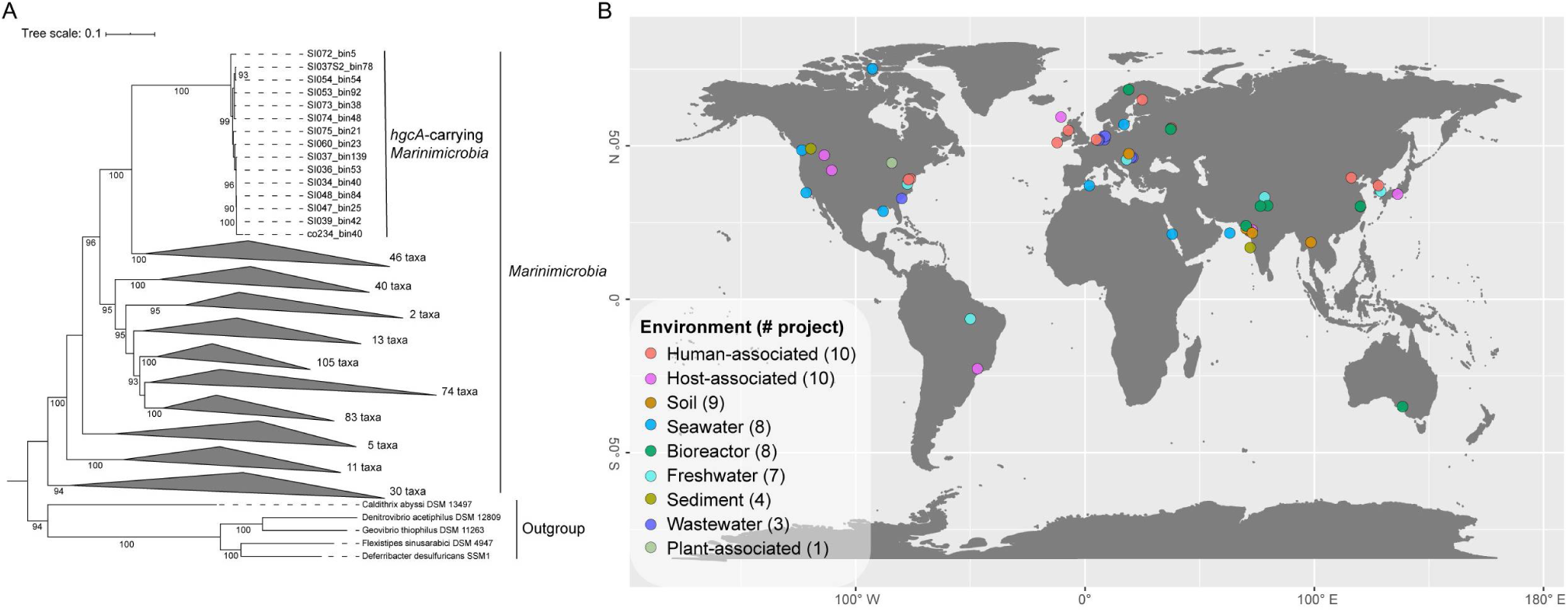
Phylogeny and global distribution of phylum *Marinimicrobia*. (A) Maximum-likelihood phylogenetic tree of phylum *Marinimicrobia* based on concatenated housekeeping genes (1,000 ultrafast bootstrap replicates; values >90% are shown at the nodes). A total of 424 *Marinimicrobia* genomes were used for the tree; *Marinimicrobia* lineages without *hgcA* genes were collapsed to simplify presentation (see Figure S6 for a more detailed tree). (B) Distribution of *hgcA*-carrying *Marinimicrobia* in various environments globally, shown as different colors; total numbers of BioProjects from each environment are shown in brackets.

### Marinimicrobia-hgcA distribution in public databases

In order to explore the global distribution of *hgcA*-carrying *Marinimicrobia*, a wider search was performed against NCBI SRA metagenomic datasets. In total, 58 BioProjects from a range of environments, including seawater, soil, sediment, freshwater, industrial wastewater, and plant rhizomes, contained *Marinimicrobia-hgcA* associated reads (Figure 4B, Table S2), suggesting a wide ecological distribution.

### Homology models of Marinimicrobia-*HgcA* and -*HgcB* proteins

As no cultivated representative of *Marinimicrobia* is currently available to assay for functional Hg methylation, we constructed homology models to test the hypothesis that HgcA and HgcB proteins encoded for by *Marinimicrobia* possess the three-dimensional structural functionality required for methylating Hg.

Analysis of the putative *Marinimicrobia*-HgcA sequence revealed a comparable domain structure (Figure 5A) to the previously characterized HgcA from the functionally validated Hg-methylating strain *Desulfovibrio desulfuricans ND132* (Parks et al 2013), including the presence of a globular domain at the N-terminus and five transmembrane spanning helices at the C-terminus. The cap-helix structure and coordinated Cys residues required for Hg(II) methylation by ND132 were conserved in the globular domain of our putative *Marinimicrobia*-HgcA (Figure S7A and Figure S7B), consistent with a conserved catalytic mechanism. The structure of ND132-HgcA was solved bound to the co-factor cobalamin required for catalytic methylation activity. The cobalamin binding pocket for ND132 was comparable to that of the *Marinimicrobia*-HgcA homology model (726.5 Å^3^ and 678.4 Å^3^, respectively), with 69% conservation of residues interacting with cobalamin. Modelling of cobalamin in the binding site of the *Marinimicrobia*-HgcA homology model revealed a similar H-bonding network for the two structures (Table S3). Modelling of the *Marinimicrobia*-HgcB protein (Figure 5C) demonstrated binding with 2[4Fe-4S] clusters at the N-terminal, similar to the structure of ND132-HgcB (Figure S7C and Figure S7D). The conserved two ferredoxin motifs (CX2CX2CX3C), and the C-terminal Cys tail in ND132-HgcB required for potentially transferring electrons and binding the Hg(II) substrate (Parks et al 2013, Rush 2018), were also observed for *Marinimicrobia*-HgcB. The structural qualities of *Marinimicrobia*-HgcA and -HgcB were further compared using Ramachandran plots (Figure 5B and 5D), with most residues located in the favored or allowed regions in both structures. Overall, this analysis strongly supports the conservation of Hg(II) methylation activity for the putative *Marinimicrobia*-HgcA and -HgcB proteins. Additionally, similar HgcAB homology models constructed for novel putative Hg methylators *Calditrichaeota* and SAR324 also support functionality (Figure S8).

**Figure 5.**
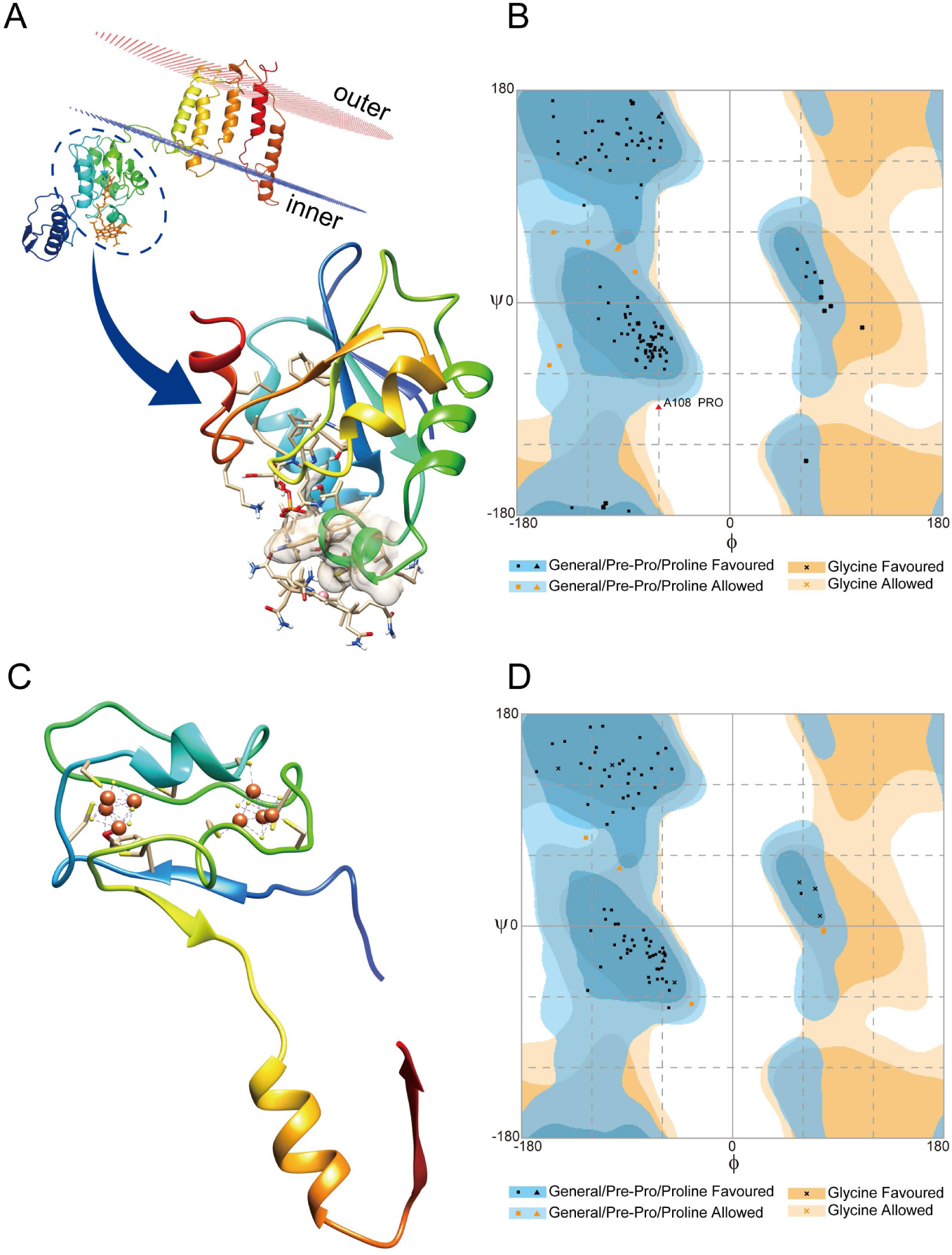
Three-dimensional homology models of *Marinimicrobia*-HgcA and -HgcB proteins. (A) Model of full-length *Marinimicrobia* HgcA, parallel to the membrane. An enlarged view of the functional domain binding to cobalamin is shown, with the interaction between cysteine and cobalt represented by a dotted line. The electrostatic surface potential of the cap-helix structure is shown. (B) Ramachandran plot of *Marinimicrobia*-HgcA model; 93.2%, 5.9%, and 0.8% of residues plotted in favoured, allowed, and outlier regions, respectively. (C) Model of full-length *Marinimicrobia*-HgcB binding with two [4Fe4S] clusters. Interactions between the protein and iron are shown by dotted lines. (D) Ramachandran plot of *Marinimicrobia*-HgcB model; 94.1%, 5.9%, and 0% of residues plotted in favoured, allowed, and outlier regions, respectively.

### Genome composition of hgcA-carrying MAGs

Metabolic pathways were inferred for MAGs by reference to Kyoto Encyclopedia of Genes and Genomes (KEGG) annotations (Figure 6). Several genes encoding for different terminal oxidases were detected, including cytochrome *c* oxidase *aa3*-type (*coxABCD*), cytochrome *c* oxidase *cbb3*-type (*ccoPQNO*), cytochrome *o* oxidase (*cyoABCD*), cytochrome *aa*_3_-600 menaquinol oxidase (*qoxABCD*), and cytochrome *bd*-type quinol oxidase (*cydAB*), demonstrating the oxygen utilization/tolerance capacities of these MAGs. Specifically, all *hgcA*-carrying *Marinimicrobia* contained cytochrome *c* oxidase *aa3*-type and cytochrome *aa*_3_-600 oxidase, and *Marinimicrobia* SI039_bin42 contained another *cbb*_3_-type oxidase. All *hgcA*-carrying SAR324, and the majority of other *hgcA*-carrying *Deltaproteobacteria*, contained cytochrome *bd* complex; and most of these also contained cytochrome *c* oxidase, and *aa3*-type and *o* ubiquinol oxidases. A few *Deltaproteobacteria* carried cytochrome *aa*_3_-600 oxidase. The two *Nitrospina* MAGs both used *cbb*_3_-type oxidase to terminate respiratory chains, and one also carried genes encoding for cytochrome *c* oxidase. In contrast, no genes involved in O_2_ respiration were observed in *hgcA*-carrying *Calditrichaeota, Verrucomicrobia, Spirochaetota*, and *Firmicutes* genomes identified from databases.

**Figure 6.**
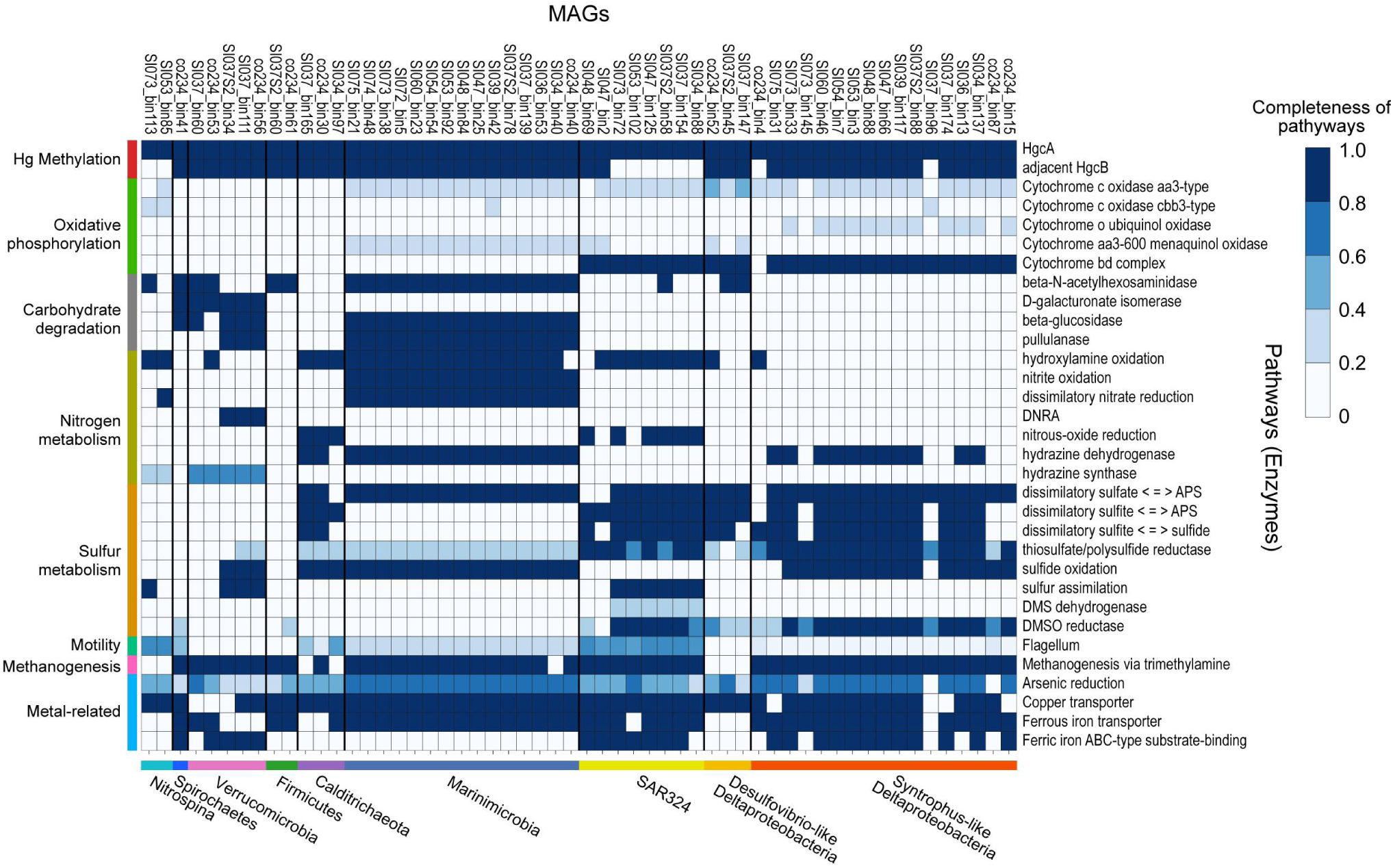
KEGG pathways of the *hgcA*-carrying MAGs. Taxonomic classifications of MAGs are represented at bottom of heatmap by different colors. Categories of pathways are represented at left side of heatmap by different colors. Color of each cell refers to completeness of enzymes involved in each pathway.

Genes involved in carbohydrate degradation pathways were found in many *hgcA*-carrying MAGs, except for *Syntrophus*-like *Deltaproteobacteria* and *Calditrichaeota*. These MAGs represent versatile metabolic capabilities in nitrogen and sulfur cycling. Notably, *Marinimicrobia* MAGs contained complete gene operons for utilising nitrogen species in electron transfers, e.g., hydroxylamine oxidoreductase (*hao*), nitrite oxidase (*nxrAB*), nitrate reductase (*narGHI*), and hydrazine dehydrogenase (*hdh*). *Deltaproteobacteria* MAGs presented various capabilities for sulfur utilisation, for which SAR324 MAGs were the only group possessing the gene *ddhA* encoding for a subunit of dimethyl sulfide (DMS) dehydrogenase. Nearly all MAGs contained some genes involved in methanogenesis and arsenic reduction, as well as transporters for Cu and Fe. Furthermore, MAGs associated with *Marinimicrobia, Calditrichaeota*, SAR324, *Spirochaetes*, and *Nitrospina* were found to carry genes encoding proteins for flagellar assembly (Figure 6).

### Novel putative marine Hg methylators in SI

In this study, SI provided a model natural ecosystem in which to study Hg methylation under oxygen-depleted conditions (Hawley et al 2017b). Our results indicate that SI water column contained zones of MeHg production, where ∼17% of total Hg was as MeHg. Similar MeHg peaks under suboxic conditions have been observed in other seawater depth profiles: the Pacific Ocean (Hammerschmidt and Bowman 2012), Arctic Ocean (Wang et al 2012), Southern Ocean (Cossa et al 2011), Arabian Sea (Chakraborty et al 2016), and Mediterranean Sea (Cossa et al 2009). Contrasting the relative abundance of potential methylators observed in the benthic zone, the observed increased abundance of *merB* genes and transcripts may provide a viable hypothesis for why MeHg levels decreased over certain depth ranges (Figure S5).

A number of known and putative Hg methylators were found, in which the phyla *Marinimicrobia* and *Calditrichaeota*, as well as the clade SAR324, are reported for the first time as carrying *hgcAB* genes (Table 1 and Figure 2). As most known Hg-methylating microorganisms derive from a limited number of phylogenetic groups (Gilmour et al 2013, Gionfriddo et al 2016, Jones et al 2019), our findings expand the database of putative microbial Hg-methylators and their microenvironments. Unfortunately, no cultivated representatives are currently available to confirm Hg methylation capability experimentally. However, protein homology modelling here provided theoretical HgcAB structures (Figure 5) that demonstrate intermolecular forces consistent with those of experimentally validated methylator, *D. desulfuricans* ND132 (Figure S7). The same methodology has been adapted to infer functionality for HgcAB proteins from *Nitrospina* (Gionfriddo et al 2016), another uncultivated potential Hg methylator (Gionfriddo et al 2016, Villar et al 2020). This approach represents a promising method for predicting the functionality of proteins encoded for by uncultivated microorganisms.

Surprisingly, *Marinimicrobia* dominated Hg-methylation capability across all water depths, and comprised a novel phylogenetic clade (Figure 4A and Figure S6). This finding supports the O_2_ adaptative capability of this clade, and both broadens and distinguishes the evolutionary history of *hgcA*. Notably, of many *hgcA* DNA sequences from the upper depths of SI seawater, only a small number were actively expressed (Figure 3C and 3D), suggesting a strong influence from oxygen level or other environmental factors. We therefore caution that abundance of *hgcA* genes, as measured by metagenomic or amplicon sequencing, cannot effectively predict the actual extent of MeHg production.

### Marinimicrobia is a widespread potential Hg methylator

The phylum *Marinimicrobia*, an uncultivated but apparently cosmopolitan group, is thought to play an important role in global biogeochemical cycling, especially across redox gradients (Bertagnolli et al 2017, Hawley et al 2017a). However, the Hg-methylation potential of *Marinimicrobia* has not previously been recognised. A few HgcA sequences from SI seawater columns were discovered by Podar et al (2015) from metagenomic data; these unidentified HgcA sequences phylogenetically co-locate with *Marinimicrobia*-HgcA identified in this study with genome resolution. Furthermore, we found *hgcA*-carrying *Marinimicrobia* exhibit a global distribution pattern (Figure 4B). Interestingly, our findings of *hgcA*-carrying *Marinimicrobia* from metagenomic datasets from Gulf of Mexico (PRJNA288120) and Canadian Arctic (PRJNA266338) waters contaminated by oil spills (Yergeau et al 2015, Yergeau et al 2017) may provide one explanation for observed *in situ* MeHg production (Liu et al 2015).

### Adaptation strategies of Hg methylators in oxygen-deficient environments

Hg methylators living in oxygen gradients may require the capability to tolerate intermittent lower levels of oxygen exposure. In this study, multiple terminal oxidases of aerobic respiratory chains were associated with several putative novel Hg methylators (Figure 6). Cytochrome *c aa33*-type oxidase, a canonical heme-copper containing terminal oxidase (Sousa et al 2012), was the most common enzyme found in *hgcA*-carrying MAGs, as well as SAR324, most *Deltaproteobacteria*, all *Marinimicrobia*, and one *Nitrospina*. Other terminal oxidase systems were also recognised; for example, genes encoding for cytochrome *c cbb*_3_-type oxidase were found in several members of *Deltaproteobacteria, Marinimicrobia*, and *Nitrospina*. This oxidase has been shown to exhibit high affinity for oxygen, enabling bacteria to respire O_2_ under both oxic and suboxic conditions (Colburn-Clifford and Allen 2010, Visser et al 2006). Cytochrome *bd* oxidase was found to be used by most *Deltaproteobacteria*; this type of oxidase has also been demonstrated to support adaptation to low-oxygen environments, because of its relative high affinity for O_2_ (Borisov et al 2011). In contrast, cytochrome *o* oxidase carried by many *Deltaproteobacteria* is an enzyme proposed to function under O_2_-rich conditions (Cotter et al 1990). As shown here, many SI Hg methylator genomes contained more than one type of terminal oxidase. Such a combination of multiple terminal oxidases has been observed for other microorganisms, facilitating electron transfer to O_2_ under variable redox potentials (Cotter et al 1990). In addition, genes encoding for flagellar proteins were identified in most SI putative Hg-methylators, enabling these bacteria to reposition optimally within redox gradients to suitable O_2_ and nutrient levels. We acknowledge, however, that cultivation experiments with *hgcAB*-bearing *Marinimicrobia* isolates are needed to elucidate their lifestyle and confirm functionality for Hg methylation.

## Materials and methods

### Mercury analysis

Seawater samples for mercury analysis were collected from SI S3 station (48°35.500 N, 123°30.300 W) in April 2010. Total and methylmercury (sum of mono- and dimethylmercury forms) determinations were made using USEPA standard methods 1631-E and 1630, respectively (USEPA 1998, USEPA 2002). In brief, total Hg was determined on filtered water samples following wet chemical oxidation by BrCl, followed by reduction by NH_2_OH and SnCl_2_, rendering all Hg species as volatile Hg(0). Hg was purged from solution by N_2_ and concentrated on a gold-coated sand cartridge, which was then heated, releasing Hg for re-concentration on a second gold cartridge for final quantification by cold-vapor atomic fluorescence spectrometry (CVAFS; Tekran 2600). Methylated Hg was first separated from the seawater matrix by KCl/H_2_SO_4_/CuSO_4_ extraction and steam distillation. Removal from seawater allowed for derivatization of MeHg into methylethyl-Hg through the use of sodium tetraethylborate, which is volatile and can be purged from solution and pre-concentrated on Tenax. The MeHg derivative was then separated from other Hg forms on a packed gas chromatography column of 15% OV-3 on Chromasorb-W at 110°C, and then rendered into Hg^0^ through pyrolysis for quantification by CVAFS. Detection limits for total and methylated Hg (3σ of reagent blanks) were 0.5 and 0.1 pM, respectively.

### Multi-omics data description

The multi-omic (metagenomic, metatranscriptomic, and metaproteomic) time-series samples from SI were taken from 2009 to 2012. A total of 84 genome-resolved metagenomic samples (Table S4) were employed in this study, including 78 samples from SI midpoint station “S3” (48°35.500 N, 123°30.300 W), 6 samples from the inlet mouth station “S4” (48°6 N 123°5 W) and the inlet end station “S2” (48°33.148 N, 123°31.969 W). A set of shotgun metatranscriptomic raw data corresponding to 36 metagenomic samples (Table S4) was used to investigate the transcriptional activity of genes of interest. Protein sequences predicted from the SI metaproteomic dataset (Hawley et al 2014) were also used to confirm the expression of target proteins in the environment. Sample collection, DNA/ RNA/ protein extraction, and sequencing methods of the datasets were described previously (Hawley et al 2017b). Briefly, seawater samples spanning six major depths (10, 100, 120, 135, 150, and 200 m) were collected and filtered onto 0.22 μm Sterivex (Millipore) filters. 1.8 ml of RNAlater (Ambion) was added to metatranscriptomic sample filters, and 1.8 ml of sucrose lysis buffer was added to metaproteomic sample filters. Filters were stored at −80 °C until processing. For metagenomic samples, Sterivex filters were thawed on ice and incubated at 37 °C for 1 h with lysozyme (Sigma). Proteinase K (Sigma) and 20% SDS were added subsequently and incubated at 55 °C for 2 h with rotation. Filters were rinsed with sucrose lysis buffer after lysate was removed. Combined lysate was extracted with phenol-chloroform followed by chloroform. The aqueous layer was washed with TE buffer (pH 8.0) for three times and concentrated to 150∼400μl. The metagenomic samples were sequenced at the DOE Joint Genome Institute (JGI) and sequenced on the Illumina HiSeq 2000 platform. For metatranscriptomic samples, total RNA was extracted using the mirVana miRNA Isolation kit (Ambion) modified for Sterivex filters. RNAlater was removed from thawed filters by extrusion and rinsed with Ringer’s solution (Sigma). After incubation at 25 °C for 20 min with rotation, Ringer’s solution was removed by extrusion. Lysozyme was added, following by incubating at 37 °C for 30 min with rotation. Lysate was removed into 15 ml tube and subjected to organic extraction. TURBO DNA-free kit (Thermofisher) was used to remove DNA and the RNeasy MinElute Cleanup kit (Qiagen) was used for purification. Metatranscriptomic shotgun libraries were generated at the JGI on the HiSeq 2000 platform. For metaproteomic samples, Bugbuster (Novagen) was added to thawed filters and incubated at 25 °C for 20∼30 min with rotation. Filters were rinsed with 1 ml lysis buffer after removing lysate. Combined lysate was then subject to buffer exchange using Amicon Ultra 10 K (Millipore) with 100 mM NH_4_HCO_3_ for three times. Urea was added to a final concentration of 8M and dithiothreitol added to a final concentration of 5 mM. Samples were incubated at 60 °C for 30 min and diluted ten times with 100mM NH_4_HCO_3_, following by digesting at 37 °C for 6 h with trypsin. C18 solid phase extraction and strong cation exchange were carried out subsequently. Tandem mass spectrometry (MS/MS) was used to sequence the protein samples, and MS-GFDB (Kim et al 2010) was used to identify peptides based on the matched SI metagenomic sequences.

### Metagenomic assembly and binning

The raw Illumina reads from metagenomic samples (2×150 bp paired-end, median 13.75 Gbp per sample) were first filtered and trimmed by Trimmomatic v3.6 (Bolger et al 2014). Samples from the same station and time point were co-assembled with MEGAHIT v1.2.8. MAGs were recovered using MetaBAT v2.14 (Kang et al 2015) and MaxBin v2.2.7 (Wu et al 2014), followed by merged and refined with MetaWRAP v1.2 (Uritskiy et al 2018). In order to further improve the quality of the MAGs, metagenomic reads were remapped to each MAG and then reassembled by SPAdes (Bankevich et al 2012) in careful mode. The qualities of derived MAGs were examined using CheckM (Parks et al 2015). 16S rRNA genes were predicted by RNAmmer (Lagesen et al 2007) and then used to assign taxonomic classifications of the MAGs. For those MAGs that lacked 16S rRNA genes, GTDB-Tk v0.3.2 (Parks et al 2018) was used to predict taxonomies according to genome contents. Genome annotation was conducted with Prokka v1.14 (Seemann 2014). The presence of *hgcAB* genes in MAGs was further confirmed by tetra-nucleotide frequencies signatures based on ESOM mapping (Ultsch and Mörchen 2005).

### Relative abundance calculation

CD-HIT v4.6 (Godzik and Li 2006) was used to choose representative *hgcA* sequences from all the MAGs under a 99% sequence identity threshold. Paired-end reads were remapped to these *hgcA* sequences with BWA-MEM algorithm (Li 2013). BBMap (http://sourceforge.net/projects/bbmap/) was used to calculate the average coverage. MicrobeCensus (Nayfach and Pollard 2015) was used to estimate the genome equivalent of every sample. The relative abundance of every representative *hgcA* in every sample was calculated by the formula “*hgcA* coverage / genome equivalent”; relative abundance of *hgcA* genes was assumed to represent that of corresponding MAGs, since *hgcA* genes are single copied in all reported methylators. To evaluate and compare expression levels of genes of interest, RPKM values were calculated for each gene, normalized for both gene length and sequencing depth..

### Search for hgcAB genes in MAGs

An in-house database of experimentally validated functional HgcAB sequences (see Table S5) was used to search for HgcAB encoding genes in MAGs by BLASTp, and BLAST results were further confirmed by examining the existence of conserved motifs ([N(V/I)WCA(A/G)(A/G)K] in HgcA and [CX2CX2CX3C] in HgcB), respectively.

### Phylogenetic analysis

HgcA sequences found in MAGs were aligned with other experimentally validated and predicted sequences (Gilmour et al 2013) by MAFFT v7 with the high-sensitivity (L-INS-i) algorithm (Standley and Katoh 2013). The alignment was trimmed by trimal v1.2 (Silla-Martínez et al 2009), followed by the maximum likelihood (ML) tree was reconstructed by IQ-TREE v1.6 (Schmidt et al 2014) under the LG+F+R9 protein substitution model chosen according to BIC. Publicly available *Marinimicrobia* 16S rRNA genes (see Table S6) were retrieved from the SILVA database (release 138; https://www.arb-silva.de/) and aligned with the 16S rRNA genes of *hgcA*-carrying *Marinimicrobia* to build an ML tree using IQ-TREE v1.6. The ML tree of all the *Marinimicrobia*-affiliated MAGs found in this study, as well as in public databases (see Table S7), was built by PhyloPhlAn2 (Segata et al 2013) using 400 universal proteins without duplication.

### Searching SRA database

To investigate the distribution of *Marinimicrobia-hgcA*, the recovered *Marinimicrobia-hgcA* gene nucleotide sequence was first employed as a query to be searched in the NCBI SRA database using the SRA search tool (Levi et al 2018), which is able to map ∼1% reads of more than 100,000 public whole shotgun (WGS) metagenomic samples to the query sequences using Bowtie2 (Langmead and Salzberg 2012). Samples that contained reads associated with the *Marinimicrobia-hgcA* gene were then checked by downloading the corresponding read set from the SRA database and mapping these reads to the *hgcA* nucleotide sequence.

### Protein homology modeling

The three-dimensional structures of the putative HgcA and HgcB sequences found in this study were built and refined using Schrödinger suit v2012 (LLC, New York, USA). The transmembrane regions were predicted by CCTOP (Dobson et al 2015). HgcAB sequences were searched against the protein data bank (PDB) using BLASTp with a word size of three. The structure of the corrinoid iron-sulfur protein (CFeSP) AcsC from *Carboxydothermus hydrogenoformans* (PDB ID: 2H9A; 1.9 Å; 28% identity and 55% similarity to *Marinimicrobia*-HgcA) and the CFeSP AcsC from *Moorella thermoacetica* (PDB ID: 4DJD; 2.38 Å; 31% identity and 59% similarity to *Marinimicrobia*-HgcA) were identified as appropriate templates for homology modeling of the HgcA N-terminus globular structured domain, according to their similarities and the resolution at which they were solved. The structure of the transmembrane bacteriorhodopsin Bop (PDB ID:2ZFE; 2.5 Å) was used to model the transmembrane region of HgcA, due to their comparable secondary structures. The homology model of HgcB proteins was built using the experimental structures of iron hydrogenase HydA from *D. desulfuricans* (PDB ID: 1HFE; 1.6 Å; 36% identity and 53% similarity to *Marinimicrobia*-HgcB) and the flavin-based caffeyl-CoA reductase CarE (PDB ID: 6FAH_A; 3.133 Å; 39% identity and 45% similarity to *Marinimicrobia*-HgcB) in a similar procedure. The stereochemical quality and accuracy of reconstructed homology models were assessed and compared to each other by generating a Ramachandran plot using the Rampage server (Lovell et al 2003) and a best model was selected. Hydrogen bonding between HgcA protein and the ligand cobalamin were predicted by Arpeggio (Jubb et al 2017), PLIP (Salentin et al 2015), Chimera (Pettersen et al 2004), and NGL Viewer (Rose and Hildebrand 2015), and consistent hydrogen bonding among the four programs was considered to be robust. Homology models for HgcAB from the Hg-methylating model strain *D. desulfuricans* ND132 were also built by the methods described above.

## Acknowledgements

H.L. was supported by a postgraduate fellowship from The University of Melbourne Environmental Microbiology Research Initiative, awarded to J.W.M. D.B.A. was supported by an Investigator Grant from the National Health and Medical Research Council (NHMRC) of Australia [GNT1174405], and by the Victorian Government OIS Program. C.H.L thanks the Woods Hole Oceanographic Institution for support and Tracy Mincer for help and inspiration. K.E.H. is supported by a Senior Medical Research Fellowship from the Viertel Foundation of Australia. H.L, D.B.H., Y.M., R.W., K.E.H and J.W.M. gratefully acknowledge the use of data generated under the auspices of the US Department of Energy (DOE) Joint Genome Institute and Office of Science User Facility, supported by the Office of Science of the U.S. Department of Energy under Contract DE-AC02-05CH11231, the G. Unger Vetlesen and Ambrose Monell Foundations, and the Natural Sciences and Engineering Research Council of Canada through grants awarded to S.J.H..

## Data availability

Data from this project have been deposited at DDBJ/ENA/GenBank under Project ID PRJNA630981. The 2088 MAGs generated by this study are available with accession numbers JABGOO000000000 to JABJQV000000000.

## Code availability

Customised scripts for conducting metagenomic and metatranscriptomic analyses are available from GitHub (https://github.com/SilentGene/MultiOmicAnalysis).

