## Supplemental Materials for "Mercury methylation by metabolically versatile and cosmopolitan marine bacteria"

Supplementary Figures:

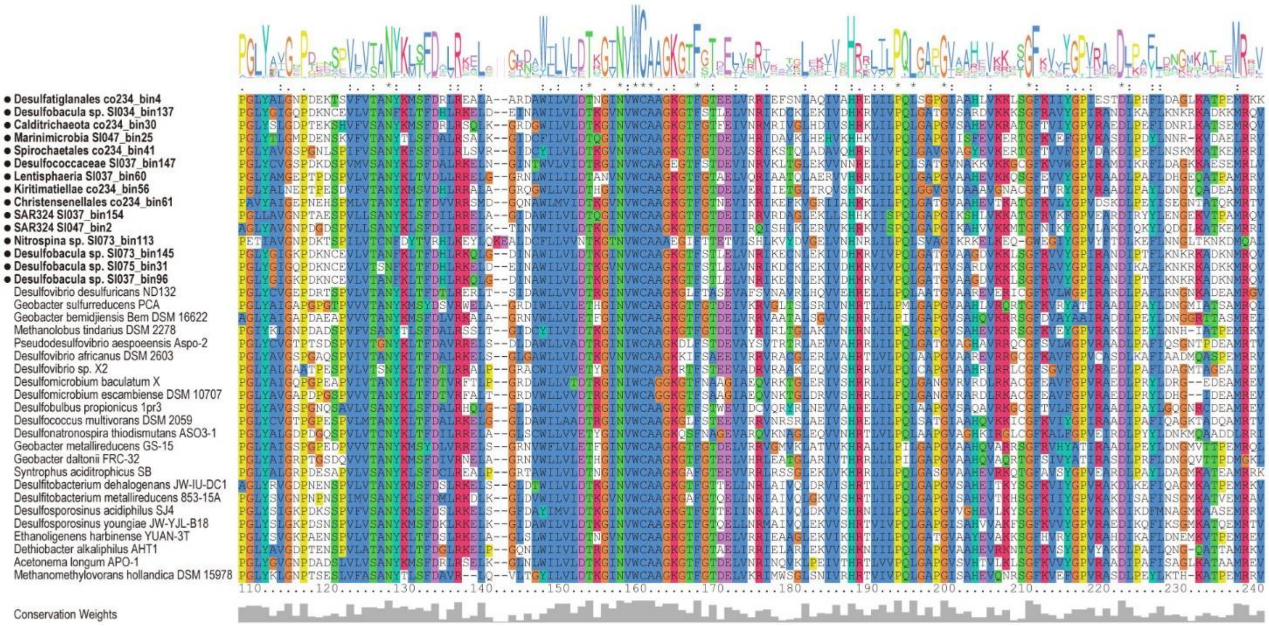


**Figure S1. Multiple *hgcA* sequence alignment.** Alignment shows 15 partial representative *hgcA* sequences found in this study (highlighted in bold and with black dots) and other experimentally confirmed *hgcA* sequences. Sequence logo above the alignment shows the occurrence of each amino acid in each position. Conservation of each sequence position is shown below the alignment.

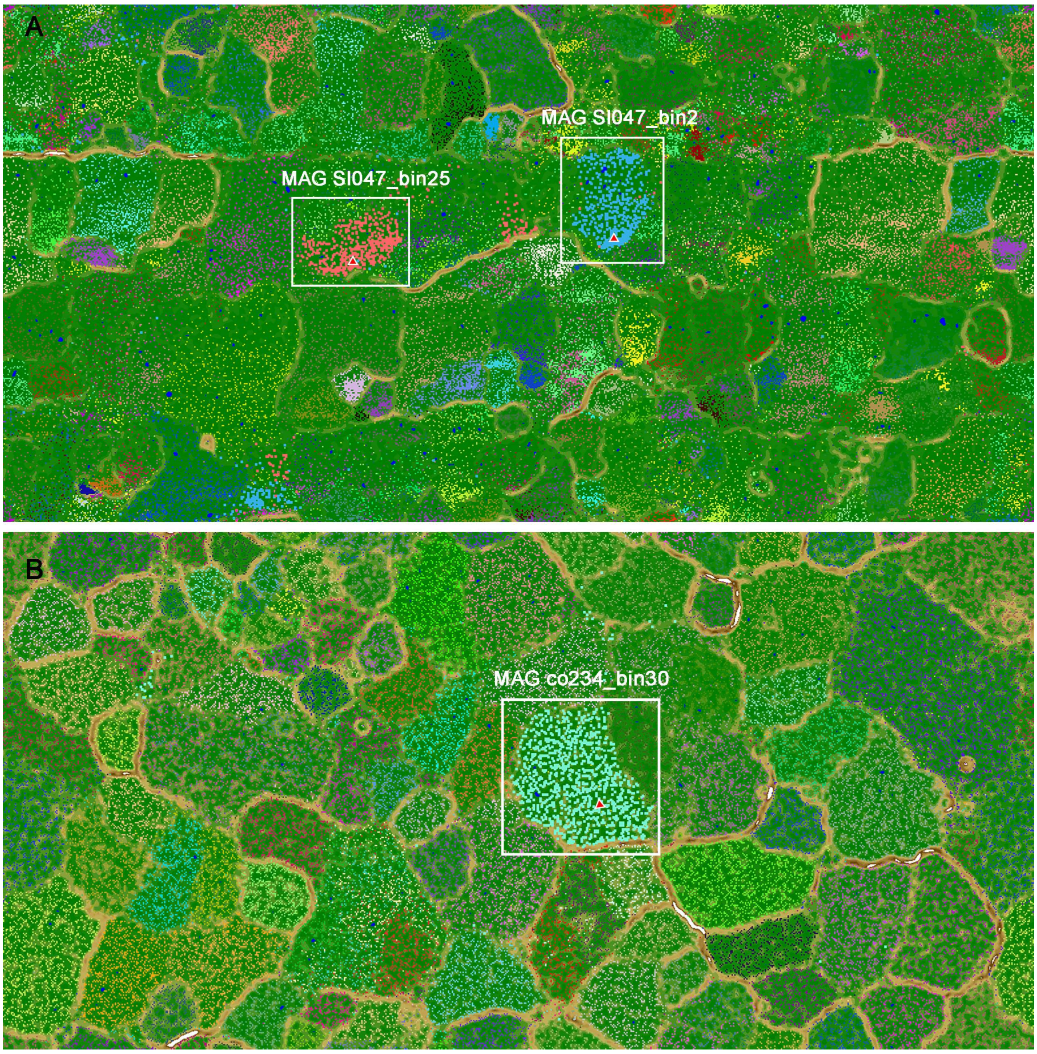


**Figure S2. Emergent self-organising maps (ESOM).** ESOMs show the three representative novel *hgcA*-carrying MAGs recovered from Saanich Inlet metagenomic datasets. Dots with different colors represent DNA fragments from different MAGs. (A) ESOM map built according to 131 MAGs recovered together with *Marinimicrobia* MAG SI047_bin25 and SAR324 MAG SI047_bin2, with the two white squares indicating DNA fragments that belonged to each of two targeted MAGs, respectively. Red triangles with white edges indicate fragments containing *hgcA* genes binned with other fragments. (B) ESOM map built according to 97 MAGs recovered together with *Calditrichaeota* MAG co234_bin30, with white square indicating DNA fragments that belonged to MAG co234_bin30. Red triangle with white edge indicates the fragment containing the *hgcA* gene.

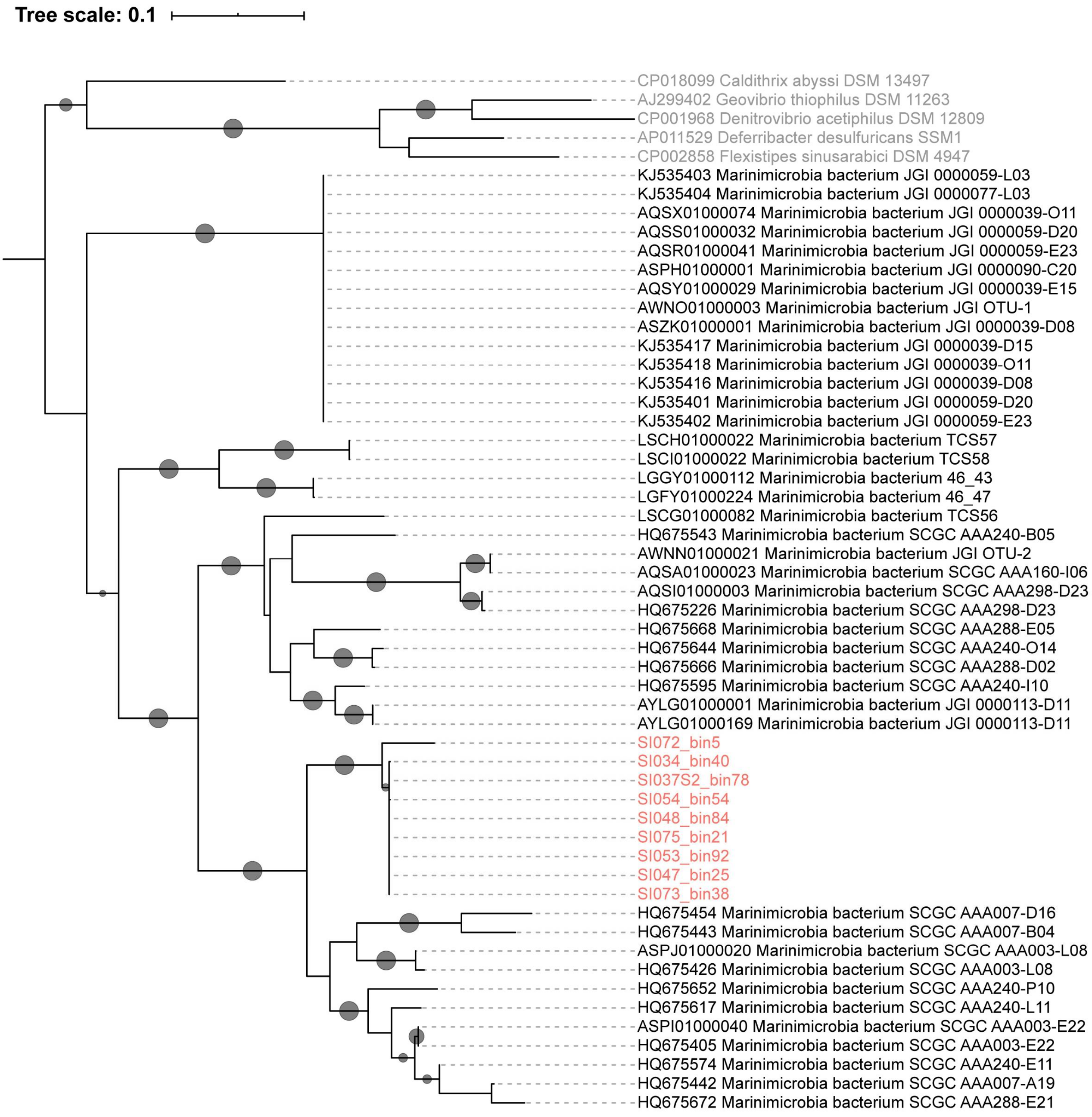


**Figure S3. Maximum-likelihood phylogenetic tree of 16S rRNA genes.** Tree was built from *hgcA*-carrying *Marinimicrobia* (red) and reference *Marinimicrobia* (black) from SILVA database. Branch support analysis was evaluated by 1000 ultrafast bootstrap replicates, and values >90% are shown by black dots at nodes.

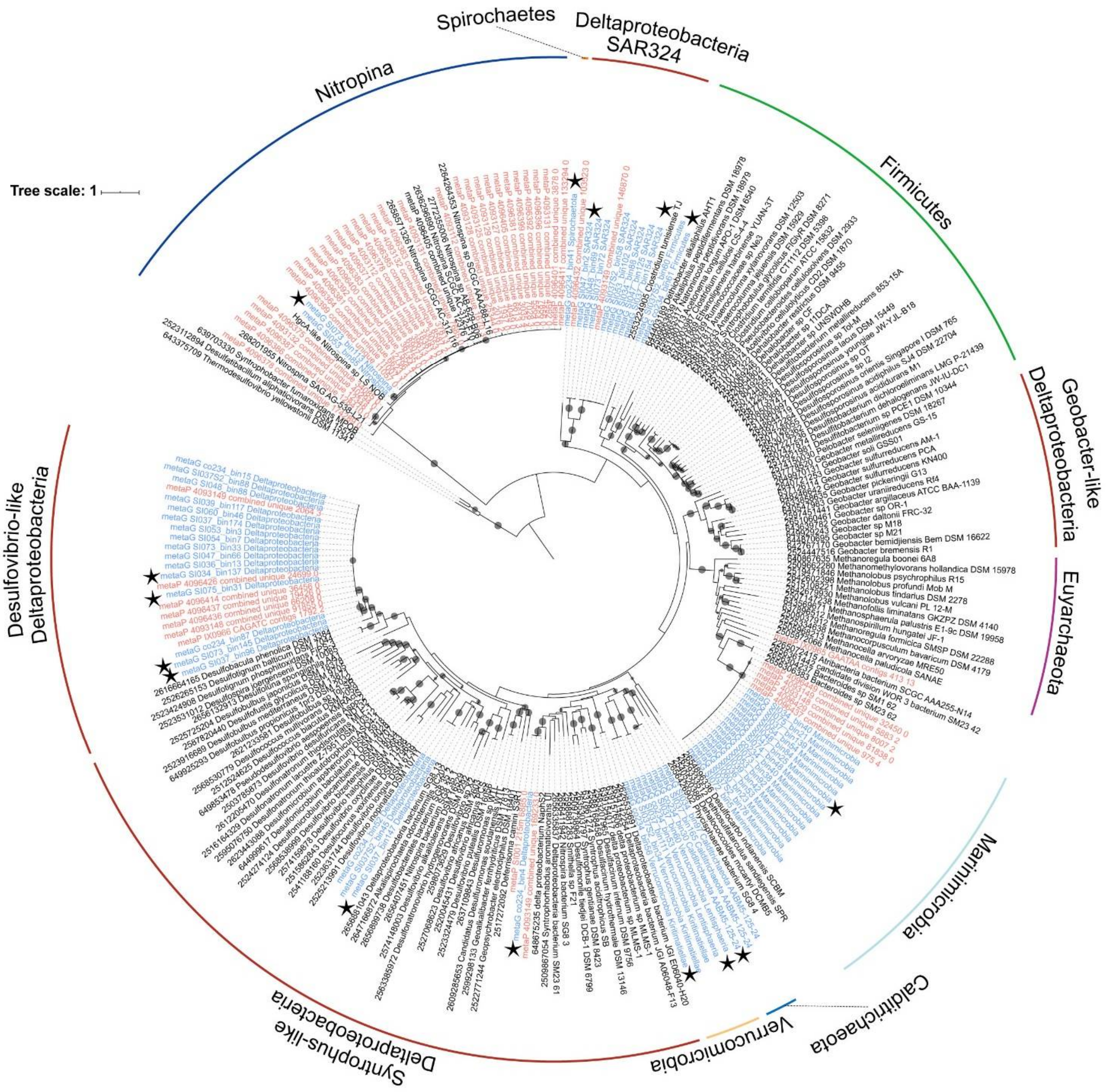


**Figure S4. Maximum-likelihood phylogenetic tree of HgcA sequences**. Tree was constructed from metagenomic (blue) and metaproteomic (red) datasets from Saanich Inlet (1,000 ultrafast bootstrap replicates; values >90% are shown by black dots at nodes). Predicted HgcA sequences from public databases and corresponding IMG gene IDs are shown in black as references. HgcA paralogues from non-methylating bacteria (shown in grey) were used as outgroups. The 15 representative HgcA sequences used in this study are indicated by stars. Taxonomic classifications of Hg methylators are labeled in the outer circle with different colors.

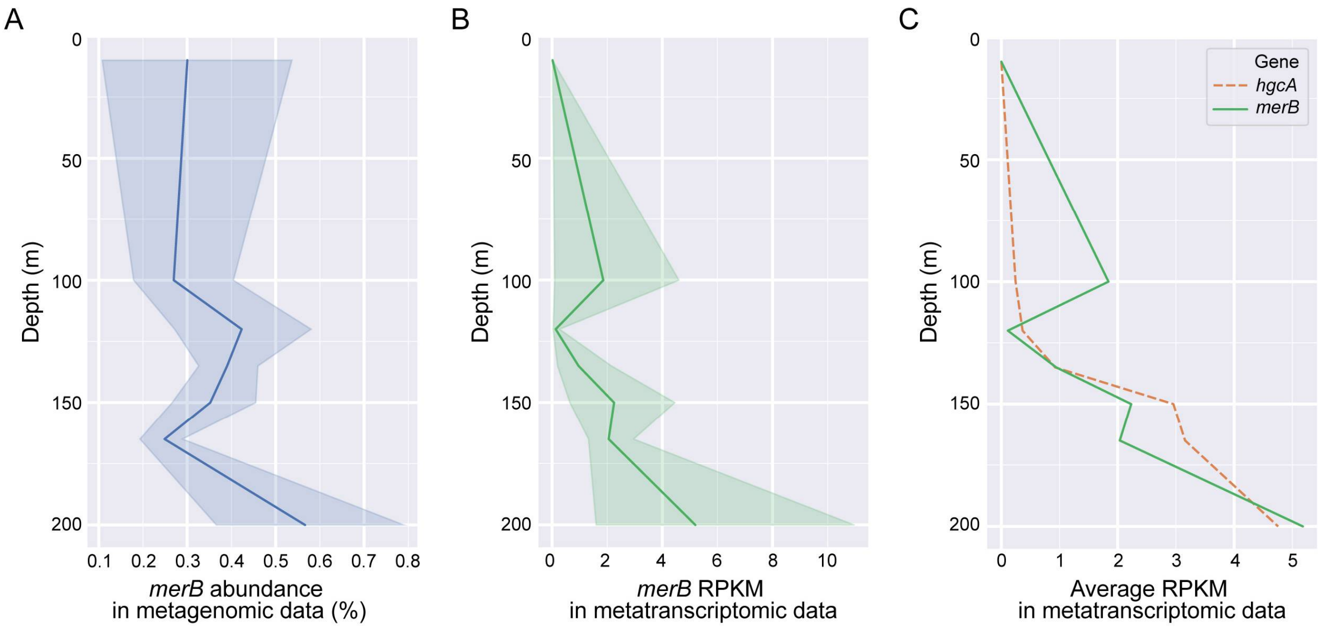


**Figure S5. DNA and RNA abundance of *merB* in Saanich Inlet datasets changing with depth.** (A) *merB* gene abundance in SI metagenomic datasets. Percentage abundance was normalized by gene length and genome equivalent for each sample. Line plotted in deep blue depicts the average abundance of *merB* with depth; light blue area depicts 95% confidence interval. (B) *merB* transcript abundance in SI metatranscriptomic datasets, as represented by RPKM values. Line plot in deep green depicts mean RPKM value of *merB* with depth; light green area depicts 95% confidence interval. (C) Comparison between RPKM values for *hgcA* (orange) and *merB* (green) in SI metatranscriptomic datasets.

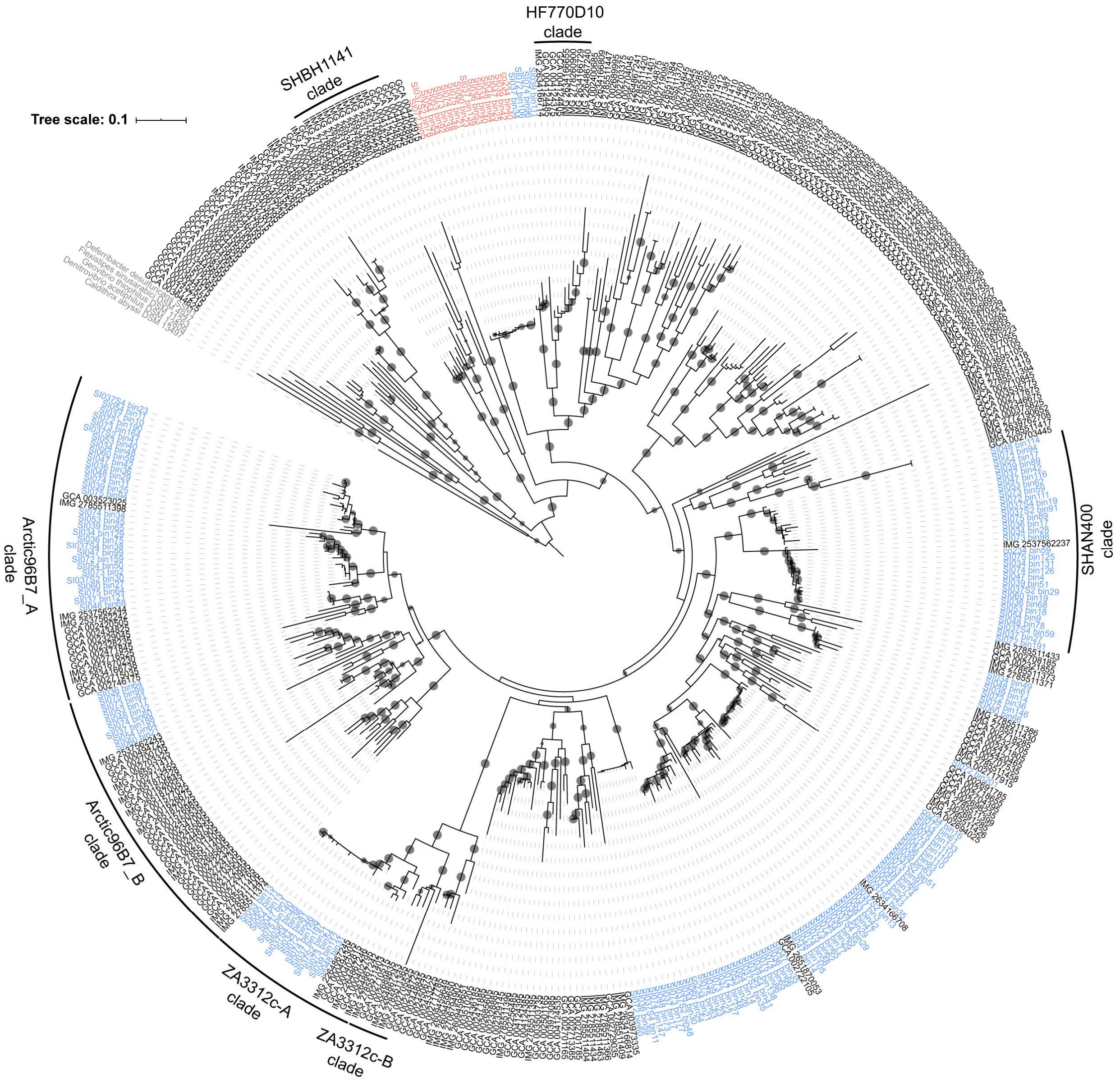


**Figure S6. Phylogenetic tree of the phylum *Marinimicrobia*.** Tree is based on concatenated housekeeping genes (1,000 ultrafast bootstrap replicates; values >90% shown by black dots at nodes). A total of 424 *Marinimicrobia* genomes were used to construct the tree. The 15 *hgcA*-carrying *Marinimicrobia* MAGs are shown in red, while other *Marinimicrobia* MAGs recovered in this study are shown in blue. *Marinimicrobia* genomes from public databases (GenBank and IMG) are shown in black, with associated accession numbers shown. Five genomes from other related phyla were used for an outgroup, and are shown in grey. *Marinimicrobia* clades as defined by Hawley et al (2017) are labeled in the outer circle.

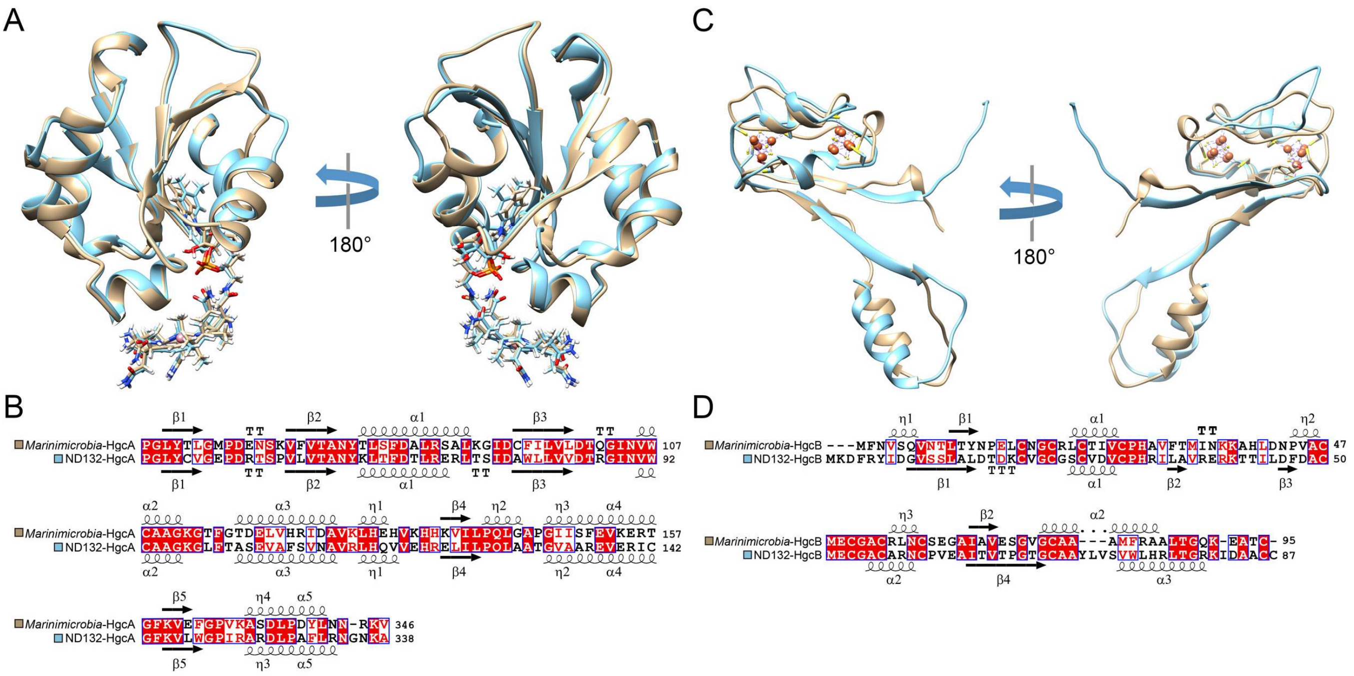


**Figure S7. Structure and sequence comparison between HgcAB from *Marinimicrobia* and *Desulfovibrio desulfuricans* ND132.** (A) Superposition of *Marinimicrobia*-HgcA (grey) and ND132-HgcA (blue) homology models. (B) Sequence alignment of *Marinimicrobia*-HgcA and ND132-HgcA amino acid sequences; secondary structure elements are indicated. A red box over a white character highlights a strict identity, a red character indicates a similarity in a group, and a blue frame indicates a similarity across groups. (C) Superposition of *Marinimicrobia*-HgcB (grey) and ND132-HgcB (blue) homology models. (D) Sequence alignment of *Marinimicrobia*-HgcB and ND132-HgcB amino acid sequences; secondary structure elements are indicated. A red box over a white character highlights a strict identity, a red character indicates a similarity in a group, and a blue frame indicates a similarity across groups.

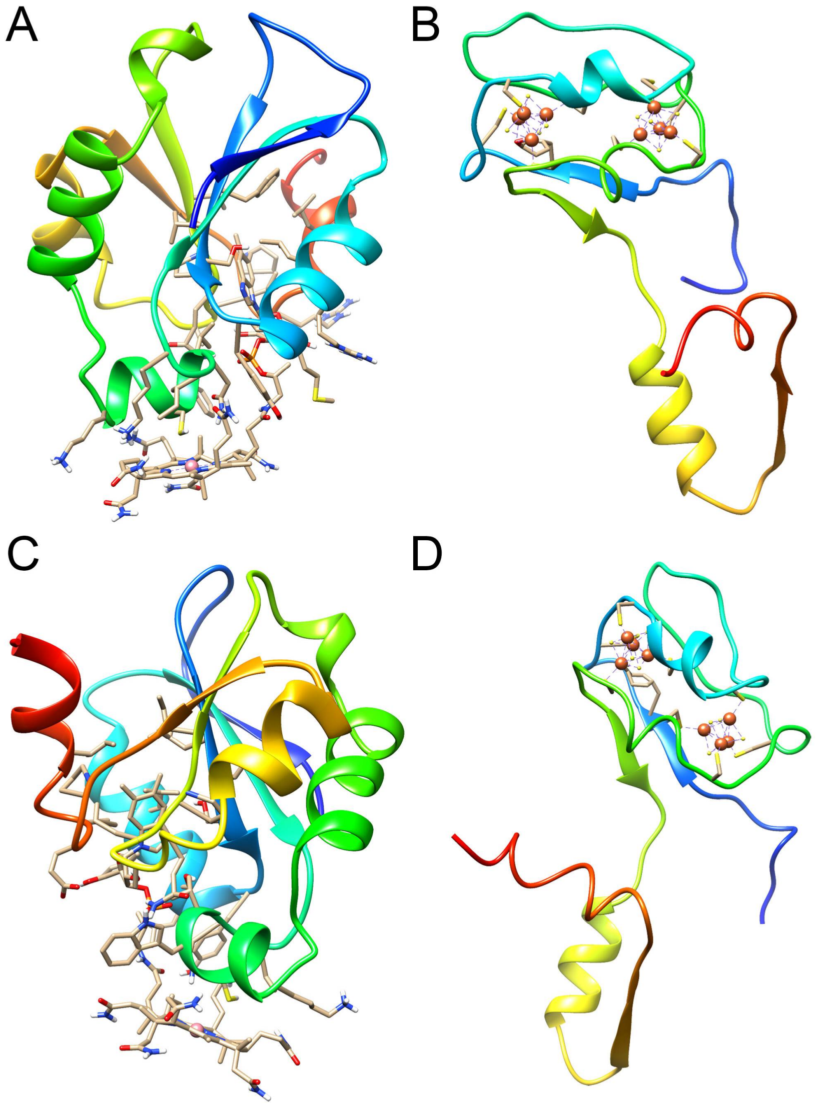


**Figure S8. Homology models of HgcA and HgcB proteins.** (A) *Calditrichaeota* HgcA model (globular domain) complexed with cobalamin. (B) *Calditrichaeota* HgcB model complexed with two [4Fe4S] ligands. (C) SAR324 HgcA model (globular domain) complexed with cobalamin. (D) SAR324 HgcB model complexed with two [4Fe4S] clusters.

Supplementary Tables:

**Table S1. Summary of all 56 MAGs recovered with *hgcA***

| MAGs | %Compl.[b] | %Conta.[b] | Size (Mbp) | # Contigs | %GC | 16S | Taxonomy |
|---|---|---|---|---|---|---|---|
| SI034_bin137 [a] | **97.58** | **2.992** | 4.94 | 304 | 40.0 | | *Desulfobacula sp.* |
| SI036_bin13 | 94.08 | **1.451** | 4.41 | 569 | 39.4 | | *Desulfobacula sp.* |
| SI037_bin174 | 91.14 | **0.161** | 4.79 | 322 | 39.9 | | *Desulfobacula sp.* |
| SI037_bin96 [a] | 71.02 | **3.87** | 2.76 | 633 | 39.5 | | *Desulfobacula sp.* |
| SI037S2_bin88 | 94.81 | **0.806** | 4.75 | 288 | 39.8 | | *Desulfobacula sp.* |
| SI039_bin117 | **97.41** | **2.096** | 5.24 | 349 | 40.0 | | *Desulfobacula sp.* |
| SI047_bin66 | **95.3** | **1.451** | 4.80 | 316 | 39.8 | | *Desulfobacula sp.* |
| SI048_bin88 | 93.22 | **0.842** | 4.98 | 332 | 39.9 | | *Desulfobacula sp.* |
| SI053_bin3 | **96.45** | **2.096** | 5.04 | 296 | 39.9 | | *Desulfobacula sp.* |
| SI054_bin7 | **97.67** | **2.096** | 5.43 | 334 | 39.9 | | *Desulfobacula sp.* |
| SI060_bin46 | **95.86** | **1.612** | 5.02 | 301 | 39.8 | | *Desulfobacula sp.* |
| SI073_bin145 [a] | 87.48 | 5.322 | 4.26 | 657 | 39.2 | | *Desulfobacula sp.* |
| SI073_bin33 | 90.96 | **3.387** | 5.21 | 425 | 40.0 | | *Desulfobacula sp.* |
| SI075_bin31[a] | 88.61 | **2.293** | 4.89 | 394 | 40.1 | | *Desulfobacula sp.* |
| co234_bin87 | 71.29 | 8.387 | 2.59 | 421 | 38.6 | | *Desulfobacula sp.* |
| co234_bin15 | 93.2 | **0.161** | 4.39 | 276 | 39.9 | | *Desulfobacula sp.* |
| co234_bin52 | 83.87 | **2.58** | 3.05 | 126 | 38.3 | | Family *Desulfococcaceae* |
| co234_bin4 [a] | 81.15 | 5.545 | 5.91 | 1000 | 50.1 | | Order *Desulfatiglandales* |
| SI037_bin147 [a] | 89.67 | **1.935** | 3.28 | 289 | 38.6 | | Order *Desulfobacterales* |
| SI037S2_bin45 | 93.51 | **2.649** | 3.24 | 215 | 38.2 | | Order *Desulfobacterales* |
| SI034_bin88 | 86.52 | 9.243 | 6.40 | 1147 | 45.3 | | SAR324 group |
| SI037_bin154 [a] | 89.02 | **3.361** | 7.85 | 512 | 45.5 | | SAR324 group |
| SI037S2_bin58 | 78.51 | **2.521** | 6.42 | 346 | 45.5 | | SAR324 group |
| SI047_bin125 | 92.38 | 5.462 | 8.98 | 616 | 45.1 | | SAR324 group |
| SI047_bin2 [a] | 74.15 | **2.941** | 3.63 | 142 | 36.2 | | SAR324 group |
| SI048_bin69 | 88.09 | 7.142 | 5.69 | 636 | 36.5 | | SAR324 group |
| SI053_bin102 | 76.25 | 8.589 | 5.60 | 1256 | 44.1 | | SAR324 group |
| SI073_bin72 | 83.07 | 5.042 | 7.92 | 781 | 45.7 | | SAR324 group |
| co234_bin40 | 78.44 | **0** | 2.78 | 52 | 37.2 | | Phylum *Marinimicrobia* |
| SI034_bin40 | **95.23** | **0** | 3.18 | 93 | 37.3 | + | Phylum *Marinimicrobia* |
| SI036_bin53 | **97.8** | **1.098** | 3.40 | 115 | 37.2 | | Phylum *Marinimicrobia* |
| SI037_bin139 | **97.8** | **0** | 3.11 | 75 | 37.4 | | Phylum *Marinimicrobia* |
| SI037S2_bin78 | **93.4** | **0** | 3.21 | 86 | 37.2 | + | Phylum *Marinimicrobia* |
| SI039_bin42 | **97.8** | **0** | 3.18 | 77 | 37.3 | | Phylum *Marinimicrobia* |
| SI047_bin25 [a] | **96.7** | **0** | 3.17 | 92 | 37.4 | + | Phylum *Marinimicrobia* |
| SI048_bin84 | 93.4 | **0** | 3.13 | 81 | 37.3 | + | Phylum *Marinimicrobia* |

| | | | | | | | |
|---|---|---|---|---|---|---|---|
| SI053_bin92 | **95.6** | **0** | 3.24 | 103 | 37.3 | + | Phylum *Marinimicrobia* |
| SI054_bin54 | **96.7** | **1.098** | 3.25 | 96 | 37.3 | + | Phylum *Marinimicrobia* |
| SI060_bin23 | 94.5 | **0** | 3.36 | 103 | 37.4 | | Phylum *Marinimicrobia* |
| SI072_bin5 | **97.8** | **1.098** | 3.65 | 144 | 37.7 | + | Phylum *Marinimicrobia* |
| SI073_bin38 | **96.7** | **0** | 3.32 | 118 | 37.5 | + | Phylum *Marinimicrobia* |
| SI074_bin48 | **97.8** | **0** | 3.38 | 113 | 37.3 | | Phylum *Marinimicrobia* |
| SI075_bin21 | **97.8** | **1.098** | 3.36 | 97 | 37.3 | + | Phylum *Marinimicrobia* |
| co234_bin30 [a] | **96.08** | **1.648** | 4.54 | 293 | 42.9 | | Phylum *Calditrichaeota* |
| SI034_bin97 | 83.8 | **4.395** | 4.68 | 804 | 42.4 | | Phylum *Calditrichaeota* |
| SI037_bin165 | 93.89 | **1.712** | 4.91 | 634 | 42.5 | | Phylum *Calditrichaeota* |
| co234_bin53 | 82.59 | 5.322 | 8.93 | 1808 | 64.2 | | Class *Lentisphaeria* |
| SI037_bin60 [a] | 83.67 | **3.914** | 8.81 | 2073 | 63.9 | | Class *Lentisphaeria* |
| co234_bin56 [a] | **95.6** | **2.027** | 4.85 | 162 | 60.6 | + | Class *Kiritimatiellae* |
| SI037_bin111 | **95.6** | **2.027** | 5.15 | 323 | 60.5 | | Class *Kiritimatiellae* |
| SI037S2_bin34 | 94.93 | **3.711** | 5.78 | 606 | 60.0 | + | Class *Kiritimatiellae* |
| co234_bin61 [a] | **98.25** | **0** | 2.67 | 97 | 47.7 | | Order *Christensenellales* |
| SI037S2_bin60 | 81.57 | **0.699** | 2.05 | 375 | 48.3 | | Order *Christensenellales* |
| co234_bin41 [a] | 72.27 | **1.253** | 4.68 | 1025 | 48.9 | | Order *Spirochaetales* |
| SI053_bin85 | 82.68 | 7.692 | 3.26 | 390 | 43.3 | | *Nitrospina sp.* |
| SI073_bin113 [a] | 74.89 | **1.709** | 2.92 | 321 | 43.1 | | *Nitrospina sp.* |

[a] Representative MAGs selected by 99% *hgcA* identity threshold

[b] Completeness above 95% and contamination below 5% are shown in bold; quality of MAGs was determined by CheckM with the “lineage_wf” pipeline.

**Table S2. Summary of NCBI BioProjects that contained *Marinimicrobia-hgcA* reads**

| BioProject ID | Study | Environment | Latitude | Longitude |
|---|---|---|---|---|
| PRJNA288120 | Gulf of Mexico water and sediments Metagenome | Seawater; Sediment | 28.66 | -88.01 |
| PRJNA245339 | Reactor Electrobiome | Bioreactor | 32.78 | -79.93 |
| PRJNA266338 | Seawater and sea ice Metagenome from the Canadian Arctic | Seawater | 75.10 | -92.64 |
| PRJNA243613 | Human stool Metagenome | Human-Associated | 55.78 | 37.62 |
| PRJNA377390 | Bombyx mori P50 Raw sequence reads | Host-Associated | 30.17 | 120.09 |
| PRJEB5982 | Microbiome from Gir Cows | Host-Associated | 22.32 | 72.97 |
| PRJNA401089 | Industrial waste Metagenome | Wastewater | 55.42 | 37.34 |
| PRJNA214608 | Anaerobic fermentation Metagenome | Bioreactor | 46.15 | 20.80 |
| PRJEB6141 | Saline desert Metagenome | Soil | 22.59 | 71.27 |
| PRJNA239997 | Anaerobic mesophilic digester fed with Spirulina | Bioreactor | 52.04 | 8.49 |
| PRJNA296946 | Gut microbiota from patients with Crohn's disease | Human-Associated | 55.45 | 37.37 |
| PRJNA189997 | Saline Desert S2 Metagenome | Soil | 22.59 | 71.27 |
| PRJEB4352 | Shotgun Sequencing of Tara Oceans DNA samples corresponding to size fractions for protist | Seawater | 37.04 | 1.90 |
| PRJNA231836 | Bioreactor enrichments from intertidal sediment | Bioreactor | 53.15 | 8.15 |
| PRJNA327439 | Discovery deep brine seawater interface metagenome sequencing | Seawater | 21.17 | 38.03 |
| PRJNA309871 | Bryoria fremontii Metatranscriptome | Host-Associated | 46.90 | -113.73 |
| PRJNA299440 | Human gut Metagenome | Human-Associated | 37.02 | 127.97 |
| PRJNA209397 | Petroleum Metagenome sequencing, Project-151 | Soil | 23.03 | 70.22 |
| PRJEB14811 | Freshwater shotgun metagenome from the surface of Nakdong River during algal bloom, Busan, South Korea | Freshwater | 35.20 | 128.99 |
| PRJNA320909 | Sludge metagenome Raw sequence reads | Wastewater | 30.26 | 120.15 |
| PRJNA255460 | Marine microorganisms cultured in chemostats | Bioreactor | 53.11 | 8.85 |
| PRJNA245804 | Microbial Diversity in an Extremely High Sulfate Lake (Spotted Lake, BC, Canada) | Freshwater; Sediment | 49.08 | -119.57 |
| PRJNA73677 | Svalbard Reindeer Rumen Metagenome | Host-Associated | 59.40 | -10.46 |
| PRJNA268268 | Geothermal Mat Community Metagenome | Sediment | 30.49 | 79.63 |
| PRJNA275775 | Human samples with multiple pathogens | Human-Associated | 39.23 | -76.17 |
| PRJNA298665 | Arabian Sea oxygen minimum zone | Seawater | 21.56 | 63.11 |
| PRJNA310400 | Soil sample Metagenome | Soil | 47.42 | 19.14 |
| PRJNA272672 | Microbial composition of soda lakes in the Carpathian Basin | Freshwater | 46.87 | 19.17 |
| PRJNA244615 | Metagenomes of Saline Desert | Soil | 23.94 | 70.19 |
| PRJNA251795 | Metagenome analysis of BTEX contaminated hypoxic groundwater | Freshwater | 45.51 | 18.17 |

| | | | | |
|---|---|---|---|---|
| PRJNA274364 | Full-scale partial-nitritation anammox (PNA) reactor Metagenome | Bioreactor | 52.05 | 6.14 |
| PRJNA296416 | AAA enrichment bioreactor | Bioreactor | 51.82 | 5.87 |
| PRJNA309469 | Marine sediment Metagenome | Sediment | 16.83 | 71.99 |
| PRJNA273799 | Baltic Sea Surface Water Metagenome | Seawater | 56.93 | 17.06 |
| PRJNA304667 | Wastewaters Metagenome | Wastewater | 30.33 | 76.49 |
| PRJNA275824 | Milk Metagenome (Jaffrabadi) | Host-Associated | 22.56 | 72.95 |
| PRJNA296849 | Neutral pH hot spring Metagenome | Freshwater | 33.23 | 78.31 |
| PRJDB5323 | Metagenomic analysis of Akoya pearl oyster (Pinctada fucata martensii) hemolymph for survey of the causative agent of Akoya oyster disease | Host-Associated | 34.20 | 136.41 |
| PRJNA385554 | The microbiomes of blowflies and houseflies | Host-Associated | -22.68 | -46.93 |
| PRJNA285742 | Anaerobe digester microbial community Metagenome | Bioreactor | 46.15 | 20.08 |
| PRJEB6070 | Potential of fecal microbiota for early stage detection of colorectal cancer | Human-Associated | 51.00 | -12.14 |
| PRJNA236764 | James River Metagenome | Freshwater | 37.53 | -77.44 |
| PRJEB8968 | Core genomes of cosmopolitan surface ocean plankton | Seawater | 34.72 | -121.54 |
| PRJNA215012 | Stordalen mire environmental genomics Metagenome | Soil | 68.35 | 19.05 |
| PRJEB5437 | Soil metagenomes profiling for forensic purposes | Soil | -35.03 | 138.57 |
| PRJNA280864 | Common bean Rust infection raw sequence reads | Plant-Associated | 44.42 | -84.30 |
| PRJEB6006 | Metagenomic approach to analyze rumen microbiome from Kankrej cows | Host-Associated | 22.31 | 72.58 |
| PRJNA298260 | Beef and dairy metagenomic samples | Host-Associated | 42.02 | -110.44 |
| PRJNA247822 | Ecological Genomics of a Seasonally Anoxic Fjord; Saanich Inlet | Seawater | 48.60 | -123.50 |
| PRJEB12449 | Reproducibility of associations between the human gut microbiome and colorectal cancer assessed in a patient population from Washington, DC, USA | Human-Associated | 39.00 | -77.00 |
| PRJEB11685 | Intestinal microbiome is related to lifetime antibiotic use in Finnish pre-school children | Human-Associated | 65.00 | 25.00 |
| PRJEB9745 | Metagenome analysis of Amlakhadi canal | Soil | 21.60 | 73.00 |
| PRJNA335944 | Lower respiratory microbes Raw sequence reads | Human-Associated | 39.56 | 116.21 |
| PRJEB12357 | Impact of faecal microbiota transplantation on the intestinal microbiome in metabolic syndrome patients | Human-Associated | 52.00 | 5.00 |
| PRJEB8347 | Temporal and technical variability of human gut metagenomes | Human-Associated | 55.02 | -7.24 |

| | | | | |
|---|---|---|---|---|
| PRJNA354598 | Brazilian copper mine Metagenome | Freshwater | -6.44 | -50.04 |
| PRJNA275825 | Milk Metagenome (Mehsani) | Host-Associated | 22.56 | 72.95 |
| PRJNA295497 | Characterisation of symbiotic cellulolytic microbial consortium enriched on napier grass | Soil | 18.54 | 98.51 |

**Table S3. Consistent hydrogen bonds of *Marinimicrobia* HgcA and *D. desulfuricans* ND132 HgcA*[*]***

| Atom1 | Atom2 |
|---|---|
| [THR] 18:A.OG1 | [B12] 500:Z.N3B |
| [B12] 500:Z.O58 | [TYR] 21:A.OH |
| [B12] 500:Z.O4 | [THR] 24:A.OG1 |
| [B12] 500:Z.O51 | [ASN] 48:A.N |
| [B12] 500:Z.O44 | [LYS] 55:A.NZ |
| [GLN] 85:A.O | [B12] 500:Z.O7R |
| [B12] 500:Z.O6R | [ALA] 111:A.N |

[*] Only hydrogen bonds connecting protein and cobalamin were predicted; bonds consistent in all four programs (Arpeggio, PLIP, NGL, and Chimera) are shown.

**Table S4. Summary of Saanich Inlet metagenomic and metatranscriptomic datasets used in this study.**

| Sample ID | Cruise ID | Year | Month | Station | Depth (m) | SRX accession (metagenomics) | SRX accession (metatranscriptomics) |
|---|---|---|---|---|---|---|---|
| SI034_S3_10 | 34 | 2009 | Jun | SI03 | 10 | SRR3724190 | - |
| SI034_S3_100 | 34 | 2009 | Jun | SI03 | 100 | SRR3724308 | - |
| SI034_S3_120 | 34 | 2009 | Jun | SI03 | 120 | SRR3724311 | - |
| SI034_S3_135 | 34 | 2009 | Jun | SI03 | 135 | SRR3724312 | - |
| SI034_S3_150 | 34 | 2009 | Jun | SI03 | 150 | SRR3724307 | - |
| SI034_S3_200 | 34 | 2009 | Jun | SI03 | 200 | SRR3724310 | - |
| | | | | | | SRR3724309 | |
| SI036_S3_100 | 36 | 2009 | Aug | SI03 | 100 | SRR3724315 | - |
| SI036_S3_120 | 36 | 2009 | Aug | SI03 | 120 | SRR3724357 | - |
| SI036_S3_135 | 36 | 2009 | Aug | SI03 | 135 | SRR3724365 | - |
| SI036_S3_150 | 36 | 2009 | Aug | SI03 | 150 | SRR3724366 | - |
| SI036_S3_200 | 36 | 2009 | Aug | SI03 | 200 | SRR3724382 | - |
| SI037_S2_100 | 37 | 2009 | Sep | SI02 | 100 | SRR3719478 | - |
| SI037_S2_150 | 37 | 2009 | Sep | SI02 | 150 | SRR3719504 | - |
| SI037_S2_200 | 37 | 2009 | Sep | SI02 | 200 | SRR3719505 | - |
| SI037_S3_10 | 37 | 2009 | Sep | SI03 | 10 | SRR3719645 | - |
| SI037_S3_100 | 37 | 2009 | Sep | SI03 | 100 | SRR3719506 | - |
| SI037_S3_110 | 37 | 2009 | Sep | SI03 | 110 | SRR3719646 | - |
| SI037_S3_120 | 37 | 2009 | Sep | SI03 | 120 | SRR3719647 | - |
| SI037_S3_125 | 37 | 2009 | Sep | SI03 | 125 | SRR3719653 | - |
| SI037_S3_130 | 37 | 2009 | Sep | SI03 | 130 | SRR3719510 | - |
| SI037_S3_150 | 37 | 2009 | Sep | SI03 | 150 | SRR3719511 | - |
| SI037_S3_200 | 37 | 2009 | Sep | SI03 | 200 | SRR3719534 | - |
| SI037_S4_100 | 37 | 2009 | Sep | SI04 | 100 | SRR3719533 | - |
| SI037_S4_130 | 37 | 2009 | Sep | SI04 | 130 | SRR3719535 | - |
| SI037_S4_150 | 37 | 2009 | Sep | SI04 | 150 | SRR3719541 | - |
| SI039_S3_10 | 39 | 2009 | Nov | SI03 | 10 | SRR3724388 | - |
| SI039_S3_100 | 39 | 2009 | Nov | SI03 | 100 | SRR3724464 | - |
| SI039_S3_120 | 39 | 2009 | Nov | SI03 | 120 | SRR3724466 | - |
| SI039_S3_135 | 39 | 2009 | Nov | SI03 | 135 | SRR3724465 | - |
| SI039_S3_150 | 39 | 2009 | Nov | SI03 | 150 | SRR3724470 | - |
| SI039_S3_200 | 39 | 2009 | Nov | SI03 | 200 | SRR3724468 | - |
| | | | | | | SRR3724467 | |
| SI047_S3_100 | 47 | 2010 | July | SI03 | 100 | SRR3724469 | SRR3719718 |
| SI047_S3_120 | 47 | 2010 | July | SI03 | 120 | SRR3724456 | SRR3719720 |
| SI047_S3_135 | 47 | 2010 | July | SI03 | 135 | SRR3724482 | SRR3719721 |
| SI047_S3_150 | 47 | 2010 | July | SI03 | 150 | SRR3724508 | SRR3719719 |

| | | | | | | | |
|---|---|---|---|---|---|---|---|
| SI047_S3_200 | 47 | 2010 | July | SI03 | 200 | SRR3724533 | SRR3719722 |
| SI048_S3_10 | 48 | 2010 | Aug | SI03 | 10 | SRR3724534 | SRR3720164 |
| SI048_S3_100 | 48 | 2010 | Aug | SI03 | 100 | SRR3724643 | SRR3720165 |
| SI048_S3_120 | 48 | 2010 | Aug | SI03 | 120 | SRR3724641 | SRR3720166 |
| SI048_S3_135 | 48 | 2010 | Aug | SI03 | 135 | SRR3724645 | SRR3720197 |
| SI048_S3_150 | 48 | 2010 | Aug | SI03 | 150 | SRR3724644 | SRR3720204 |
| SI048_S3_200 | 48 | 2010 | Aug | SI03 | 200 | SRR3724646 | SRR3720199 |
| SI053_S3_10 | 53 | 2011 | Jan | SI03 | 10 | SRR3724566 | - |
| SI053_S3_100 | 53 | 2011 | Jan | SI03 | 100 | SRR3724647 | - |
| SI053_S3_120 | 53 | 2011 | Jan | SI03 | 120 | SRR3724648 | - |
| SI053_S3_135 | 53 | 2011 | Jan | SI03 | 135 | SRR3725015 | - |
| SI053_S3_150 | 53 | 2011 | Jan | SI03 | 150 | SRR3725354 | - |
| SI053_S3_200 | 53 | 2011 | Jan | SI03 | 200 | SRR3725355 | - |
| SI054_S3_100 | 54 | 2011 | Feb | SI03 | 100 | SRR3725054 | SRR3720205 |
| SI054_S3_120 | 54 | 2011 | Feb | SI03 | 120 | SRR3725353 | SRR3720202 |
| SI054_S3_135 | 54 | 2011 | Feb | SI03 | 135 | SRR3725726 | SRR3720203 |
| SI054_S3_150 | 54 | 2011 | Feb | SI03 | 150 | SRR3725727 | SRR3720200 |
| SI054_S3_200 | 54 | 2011 | Feb | SI03 | 200 | SRR3725729 | SRR3720198 |
| SI060_S3_100 | 60 | 2011 | Aug | SI03 | 100 | SRR3725728 | - |
| SI060_S3_150 | 60 | 2011 | Aug | SI03 | 150 | SRR3725730 | - |
| SI060_S3_200 | 60 | 2011 | Aug | SI03 | 200 | SRR3725731 | - |
| SI072_S3_10 | 72 | 2012 | Aug | SI03 | 10 | SRR3719539 | SRR3719270 |
| SI072_S3_100 | 72 | 2012 | Aug | SI03 | 100 | SRR3719544 | SRR3719269<br>SRR3719279 |
| SI072_S3_120 | 72 | 2012 | Aug | SI03 | 120 | SRR3719545 | - |
| SI072_S3_135 | 72 | 2012 | Aug | SI03 | 135 | SRR3719563 | SRR3719361<br>SRR3719367 |
| SI072_S3_150 | 72 | 2012 | Aug | SI03 | 150 | SRR3719562 | SRR3719365<br>SRR3719369 |
| SI072_S3_165 | 72 | 2012 | Aug | SI03 | 165 | SRR3719654 | SRR3719366<br>SRR3719364 |
| SI072_S3_200 | 72 | 2012 | Aug | SI03 | 200 | SRR3719564 | SRR3719350<br>SRR3719363 |
| SI073_S3_10 | 73 | 2012 | Aug | SI03 | 10 | SRR3719565 | SRR3719362 |
| SI073_S3_100 | 73 | 2012 | Aug | SI03 | 100 | SRR3719566 | - |
| SI073_S3_120 | 73 | 2012 | Aug | SI03 | 120 | SRR3719567 | - |
| SI073_S3_135 | 73 | 2012 | Aug | SI03 | 135 | SRR3719568 | - |
| SI073_S3_150 | 73 | 2012 | Aug | SI03 | 150 | SRR3719569 | - |
| SI073_S3_165 | 73 | 2012 | Aug | SI03 | 165 | SRR3719655 | SRR3719368<br>SRR3719398 |

| | | | | | | | |
|---|---|---|---|---|---|---|---|
| SI073_S3_200 | 73 | 2012 | Aug | SI03 | 200 | SRR3719571 | - |
| SI074_S3_10 | 74 | 2012 | Sep | SI03 | 10 | SRR3719572 | SRR3719405 |
| SI074_S3_100 | 74 | 2012 | Sep | SI03 | 100 | SRR3719573 | SRR3719400 |
| | | | | | | | SRR3719406 |
| SI074_S3_120 | 74 | 2012 | Sep | SI03 | 120 | SRR3719574 | SRR3719402 |
| SI074_S3_135 | 74 | 2012 | Sep | SI03 | 135 | SRR3719625 | SRR3719399 |
| | | | | | | | SRR3719404 |
| SI074_S3_150 | 74 | 2012 | Sep | SI03 | 150 | SRR3719629 | SRR3719403 |
| | | | | | | | SRR3719407 |
| SI074_S3_165 | 74 | 2012 | Sep | SI03 | 165 | SRR3719657 | SRR3719408 |
| | | | | | | | SRR3719438 |
| SI074_S3_200 | 74 | 2012 | Sep | SI03 | 200 | SRR3719644 | SRR3719437 |
| | | | | | | | SRR3719439 |
| SI075_S3_10 | 75 | 2012 | Sep | SI03 | 10 | SRR3719656 | - |
| SI075_S3_100 | 75 | 2012 | Sep | SI03 | 100 | SRR3719691 | SRR3719447 |
| | | | | | | | SRR3719451 |
| SI075_S3_120 | 75 | 2012 | Sep | SI03 | 120 | SRR3719692 | SRR3719449 |
| | | | | | | | SRR3719448 |
| SI075_S3_135 | 75 | 2012 | Sep | SI03 | 135 | SRR3719693 | SRR3719450 |
| | | | | | | | SRR3719465 |
| SI075_S3_150 | 75 | 2012 | Sep | SI03 | 135 | SRR3719696 | SRR3719463 |
| | | | | | | | SRR3719464 |
| SI075_S3_165 | 75 | 2012 | Sep | SI03 | 165 | SRR3719695 | - |
| SI075_S3_200 | 75 | 2012 | Sep | SI03 | 200 | SRR3719708 | SRR3719475 |
| | | | | | | | SRR3719477 |

**Table S5. Experimentally validated Hg methylators and their *hgcAB* gene IDs in IMG database used in this study.**

| Genome name | *hgcA* gene ID | *hgcB* gene ID |
|---|---|---|
| *Acetonema longum* APO-1 | 651580713 | 651580712 |
| *Desulfitobacterium dehalogenans* JW-IU-DC1 | 2507427874 | 2507427873 |
| *Desulfitobacterium metallireducens* 853-15A | 2507244055 | 2507244054 |
| *Desulfobulbus propionicus* 1pr3 | 649925293 | 649925292 |
| *Desulfococcus multivorans* DSM 2059 | 2568530779 | 2568530778 |
| *Desulfomicrobium baculatum* DSM 4028 | 644999617 | 644999616 |
| *Desulfomicrobium escambiense* DSM 10707 | 2524274124 | 2524274123 |
| *Desulfonatronospira thiodismutans* ASO3-1 | 644405450 | 644405451 |
| *Desulfosporosinus acidiphilus* SJ4 | 2507126055 | 2507126056 |
| *Desulfosporosinus youngiae* JW-YJL-B18 | 2508509747 | 2508509746 |
| *Desulfovibrio africanus* DSM 2603 | 2527068623 | 2527068625 |
| *Desulfovibrio desulfuricans* ND132 | 2503785873 | 2503785874 |
| *Desulfovibrio sp.* X2 | 2598073629 | 2598073630 |
| *Dethiobacter alkaliphilus* AHT1 | 644403189 | 644403188 |
| *Ethanoligenens harbinense* YUAN-3T | 649821869 | 649821870 |
| *Geobacter bemidjiensis* Bem | 642767170 | 642767171 |
| *Geobacter daltonii* FRC-32 | 643639782 | 643639781 |
| *Geobacter metallireducens* GS-15 | 637778520 | 637778521 |
| *Geobacter sulfurreducens* PCA | 637126114 | 637126115 |
| *Methanolobus tindarius* DSM 2278 | 2515108221 | 2515108220 |
| *Methanomethylovorans hollandica* DSM 15978 | 2509662280 | 2509662279 |
| *Pseudodesulfovibrio aespoeensis* Aspo-2 | 649853478 | 649853479 |
| *Syntrophus aciditrophicus* SB | 637861274 | 637861273 |

**Table S6 *Marinimicrobia* 16S rRNA gene accession numbers from the SILVA database.**

| SILVA accession |
|---|
| AQSA01000023 |
| AQSI01000003 |
| AQSR01000041 |
| AQSS01000032 |
| AQSX01000074 |
| AQSY01000029 |
| ASPH01000001 |
| ASPI01000040 |
| ASPJ01000020 |
| ASZK01000001 |
| AWNN01000021 |
| AWNO01000003 |
| AYLG01000001 |
| AYLG01000169 |
| HQ675226 |
| HQ675405 |
| HQ675426 |
| HQ675442 |
| HQ675443 |
| HQ675454 |
| HQ675543 |
| HQ675574 |
| HQ675595 |
| HQ675617 |
| HQ675644 |
| HQ675652 |
| HQ675666 |
| HQ675668 |
| HQ675672 |
| KJ535401 |
| KJ535402 |
| KJ535403 |
| KJ535404 |
| KJ535416 |
| KJ535417 |
| KJ535418 |
| LGFY01000224 |
| LGGY01000112 |

| |
|---|
| LSCG01000082 |
| LSCH01000022 |
| LSCI01000022 |

**Table S7. Publicly available *Marinimicrobia* genomes used in this study.**

| | |
|---|---|
| **IMG genome ID** | 2264867236, 2264867237, 2264867238, 2264867239, 2264867240, 2264867241, 2264867242, 2264867243, 2264867245, 2264867246, 2527291526, 2537562237, 2537562238, 2537562239, 2537562240, 2537562241, 2537562242, 2537562243, 2537562244, 2609459649, 2609459650, 2609459651, 2634166655, 2634166708, 2634166714, 2634166729, 2634166738, 2634166809, 2634166814, 2639762708, 2651870051, 2651870052, 2651870053, 2651870223, 2651870224, 2651870225, 2651870226, 2651870227, 2651870228, 2651870229, 2651870230, 2693429801, 2693429802, 2706794902, 2772190652, 2778260900, 2785511365, 2785511366, 2785511370, 2785511371, 2785511373, 2785511374, 2785511376, 2785511378, 2785511382, 2785511384, 2785511386, 2785511387, 2785511394, 2785511395, 2785511396, 2785511398, 2785511399, 2785511401, 2785511403, 2785511404, 2785511405, 2785511406, 2785511408, 2785511409, 2785511410, 2785511411, 2785511412, 2785511417, 2785511418, 2785511419, 2785511420, 2785511426, 2785511430, 2785511432, 2785511433, 2785511434, 2785511437, 2785511440, 2785511443, 2785511445, 2785511447, 2785511458, 2785511460, 2785511462, 2785511463, 2786546459, 2786546460, 2786546461, 2786546462 |
| **GenBank assembly accession** | GCA_000375825.1, GCA_000398105.1, GCA_000402135.1, GCA_000402455.1, GCA_000402795.1, GCA_000402815.1, GCA_000403015.1, GCA_000403095.1, GCA_000405265.1, GCA_000405825.1, GCA_000494305.1, GCA_000494725.1, GCA_001508355.1, GCA_001509335.1, GCA_001576965.1, GCA_001576985.1, GCA_001577025.1, GCA_001577055.1, GCA_001577065.1, GCA_001577105.1, GCA_001577215.1, GCA_001626655.1, GCA_001626675.1, GCA_001626685.1, GCA_001626695.1, GCA_001626725.1, GCA_001626755.1, GCA_001626845.1, GCA_002172545.1, GCA_002256485.1, GCA_002311275.1, GCA_002311395.1, GCA_002311975.1, GCA_002321855.1, GCA_002327705.1, GCA_002327805.1, GCA_002328235.1, GCA_002328255.1, GCA_002328285.1, GCA_002328325.1, GCA_002328885.1, GCA_002328905.1, GCA_002329045.1, GCA_002329065.1, GCA_002329125.1, GCA_002329145.1, GCA_002346675.1, GCA_002346965.1, GCA_002347005.1, GCA_002347855.1, GCA_002348725.1, GCA_002365365.1, GCA_002400685.1, GCA_002402135.1, GCA_002403015.1, GCA_002420465.1, GCA_002420505.1, GCA_002420565.1, GCA_002420645.1, GCA_002433225.1, GCA_002434825.1, GCA_002434835.1, GCA_002434865.1, GCA_002436185.1, GCA_002436235.1, GCA_002448755.1, GCA_002448795.1, GCA_002448845.1, GCA_002450535.1, GCA_002450675.1, GCA_002452555.1, GCA_002455835.1, GCA_002471705.1, GCA_002471805.1, GCA_002471845.1, GCA_002471865.1, GCA_002471885.1, GCA_002501165.1, GCA_002591605.1, GCA_002685605.1, GCA_002686395.1, GCA_002689835.1, GCA_002689995.1, GCA_002691285.1, GCA_002691365.1, GCA_002691785.1, GCA_002693365.1, GCA_002694025.1, GCA_002695285.1, GCA_002695845.1, GCA_002698235.1, GCA_002700705.1, GCA_002701165.1, GCA_002701785.1, GCA_002703325.1, GCA_002703375.1, GCA_002703445.1, GCA_002703505.1, GCA_002703645.1, GCA_002704045.1, GCA_002704395.1, GCA_002704685.1, GCA_002704845.1, GCA_002704875.1, GCA_002704905.1, GCA_002705525.1, GCA_002705605.1, GCA_002707535.1, GCA_002707645.1, GCA_002707765.1, GCA_002707905.1, GCA_002708185.1, GCA_002710515.1, GCA_002710845.1, GCA_002711065.1, GCA_002713385.1, GCA_002713455.1, GCA_002715035.1, GCA_002715465.1, GCA_002716525.1, GCA_002717695.1, GCA_002717765.1, GCA_002717785.1, GCA_002717875.1, GCA_002717915.1, |

| | |
|---|---|
| | GCA_002717995.1, GCA_002718285.1, GCA_002718585.1, GCA_002718655.1, GCA_002719095.1, GCA_002719835.1, GCA_002720995.1, GCA_002722105.1, GCA_002722325.1, GCA_002724735.1, GCA_002729035.1, GCA_002729175.1, GCA_002731745.1, GCA_002746175.1, GCA_003445695.1, GCA_003503735.1, GCA_003510205.1, GCA_003512065.1, GCA_003523025.1, GCA_003527965.1, GCA_003535775.1, GCA_003538075.1, GCA_003644565.1, GCA_003644575.1, GCA_003644595.1, GCA_003644625.1, GCA_003645825.1, GCA_003645835.1, GCA_003645845.1, GCA_003695105.1, GCA_003973085.1, GCA_003973335.1, GCA_003973365.1, GCA_004124295.1, GCA_004124375.1, GCA_004124385.1, GCA_004124395.1, GCA_004124405.1, GCA_004124455.1, GCA_004124465.1, GCA_004124475.1, GCA_004124485.1, GCA_004124535.1, GCA_004124875.1, GCA_004402895.1, GCA_004402915.1, GCA_004402925.1 |
